# Epigenetic control of coordinated hematopoietic and cardiovascular development by Rnf2 in zebrafish

**DOI:** 10.1101/2020.12.14.422669

**Authors:** XX Peng, G Feng, YH Sun

## Abstract

Early embryogenesis requires the coordinated development of cardiovascular and hematopoietic lineages. However, the underlying cellular and genetic mechanisms are poorly understood. Here, we show that Rnf2, the core enzymatic component of Polycomb repressive complex 1 (PRC1), plays an important role in the control of cardiovascular and hematopoietic development and differentiation via suppressing the master hematoendothelial progenitor genes in zebrafish. In the absence of Rnf2, a group of transcription factor (TF) genes crucial for hematoendothelial specification such as *etv2*, *gata2*, *lmo2* and *tal1* are significantly up-regulated, which causes an expansion of hematopoietic and endothelial progenitors at the expense of myocardial differentiation, resulting in severe defects in both cardiogenesis and hematopoiesis. Although the number of hematopoietic stem cells (HSCs) is increased, both primitive and definitive waves of hematopoiesis are severely compromised in *rnf2* mutant embryos, suggesting that Rnf2 is required for differentiation of blood progenitor cells. Combined ChIP-seq and RNA-seq analysis shows that Rnf2 directly binds to key hematoendothelial progenitor genes and represses its expression. We further show that Rnf2-mediated gene repression depends on its H2Aub1 catalytic activity. We propose that PRC1/Rnf2-mediated epigenetic mechanism plays a key role in coordinated development of cardiovascular and hematopoietic lineages by repressing key hematoendothelial progenitor genes.

**Highlights:**

1. Rnf2 is required for suppressing the expression of key hematoendothelial TF genes in precursors and its differentiated descendants.
2. Rnf2 mutant zebrafish embryos display defective hematopoiesis and cardiogenesis.
3. Loss of Rnf2 results in increased HSC numbers and arrested differentiation, hallmarks of leukemia.
4. Rnf2 suppresses hematoendothelial progenitor genes via depositing H2Aub1.

## Introduction

There is a close relationship between cardiovascular and hematopoietic lineages throughout embryogenesis (Nakano et al., 2013). During gastrulation, embryonic myocardial, endocardial and hematoendothelial lineages may originate from mesodermal precursors called multipotential cardiovascular progenitors (MCPs) (Misfeldt et al., 2009; Milgrom-Hoffman et al., 2011; Saint-Jean et al., 2019). Soon after gastrulation, myocardial and hematoendothelial cells develop in close proximity at the anterior lateral plate mesoderm (ALPM) and are mutually antagonistic which is likely established by antagonizing transcription factors expressed in the respective cell lineages (Tu and Chi, 2012; Palencia-Desai et al., 2011). Hematoendothelial/endocardial and myocardial progenitor populations are detectable by expression of transcription factors (TFs) Gata2/Lmo2/Tal1/Etv2 and Nkx2.5/Hand2, respectively (Prummel et al., 2020). Overexpression of either *gata2* or *etv2* (or *tal1*) in hemangioblasts or ALPM promotes hematopoietic development and inhibits cardiac differentiation (Castano et al., 2019; Chagraoui et al., 2020; Van Handel et al., 2012), whereas loss of gene function leads to ectopic induction of myocardial cells at the expense of blood and endothelial cells (Chestnut et al., 2020; Gering et al., 2003; Schoenebeck et al., 2007). Analogously, overexpression of *nkx2.5/hand2* prohibits hematopoiesis and vasculogenesis (Yin et al., 2020). Interestingly, endocardium can contribute to transient definitive hematopoiesis, and endocardial haematopoiesis depends on cardiac specification TFs such as Nkx2.5 and Isl1 (Nakano et al., 2013). Thus, proper expression of master hematoendothelial and myocardial TFs is critical for the coordinated development of cardiovascular and hematopoietic lineages. Consistently, cardiovascular and hematopoietic abnormalities simultaneously occur in human diseases such as Down syndrome and Digeorge syndrome and that master transcriptional regulators of primitive hematopoiesis are closely involved in cardiovascular development and diseases (Yin et al., 2020). Therefore, it is of great importance to understand the coordinated development of cardiovascular and hematopoietic lineages. However, the underlying cellular and genetic mechanisms are poorly understood.

Polycomb group proteins (PcGs) are important epigenetic repressors that play important roles in embryonic stem cell (ESC) and hematopoietic stem cell (HSC) regulation, cell fate determination, cell lineage restriction and organogenesis (Blackledge et al., 2014; Chagraoui et al., 2020; Di Carlo et al., 2018; He et al., 2011; Schuettengruber et al., 2017), and abnormal PcG expression and mutations in PcG genes are frequently associated with various types of hematopoietic neoplasms and cardiovascular diseases (Radulovic et al., 2013; Wang, 2012). PcGs are mainly consisted of two complexes: PRC1 and PRC2. PRC2 places the repressive H3K27me3 mark via its EZH2 protein subunit, and PRC1 catalyzes monoubiquitination histone H2A lysine119 (H2Aub1) through the E3 ligase RNF2/Ring1 (Gao et al., 2012). The two PcG complexes (PRC1/PRC2) may work in concert to silence the developmental genes while keeping them in a poised state, and its loss of function leads to de-repression of developmental genes, resulting in loss of ESC stemness or various developmental defects in animals (Endoh et al., 2012; Illingworth et al., 2015; Sun et al., 2020; Wang et al., 2012). PRC1/RNF2 have been shown to stabilize the expression of developmental genes during embryogenesis, organogenesis and homeostatic development of adult tissues (Chrispijn et al., 2019; Ohtsubo et al., 2008; Van der Velden et al., 2011; Vocken et al., 2003). However, whether and how PRC1/Rnf2 are involved in the control of coordinated hematopoietic and cardiovascular lineage development remain elusive.

In this work, we hypothesize that Rnf2/PRC1-mediated epigenetic mechanism plays important roles in cardiovascular and hematopoietic lineage development and differentiation during zebrafish embryogenesis. We show that Rnf2 is required for coordinated development of cardiovascular and hematopoietic lineages by suppressing the expression of key hematoendothelial progenitor genes, in precursors at the ALPM and its differentiated descendants such as heart and blood cells. Mechanistically, Rnf2 directly binds to master hematoendothelial progenitor genes and represses its expression by depositing H2Aub1, and its loss leads to severe defects in cardiogenesis and hematopoiesis.

## Materials and methods

### Zebrafish lines and maintenance

Zebrafish were raised and maintained according to standard laboratory procedures. Wild-type AB line and transgenic line Tg (*myl7*: EGFP) were used in this study. All experiment procedures involving animals are approved by Institutional Animal Care and Use committee at the Institute of Hydrobiology, Chinese Academy of Sciences, Wuhan, China.

### Generation of *rnf2* mutants

Zebrafish *rnf2* mutants were generated by using CRISPR/Cas9 technology. gRNA was designed targeting the exon 3 of the *rnf2* gene. Embryos at the 1-cell stage were co-injected with 200 ng/µL Cas9 mRNA and 80 ng/µL guide RNA. The genomic DNA of 20 injected embryos at 24 hpf was extracted and subjected to PCR amplification. A DNA fragment containing the *rnf2* target site was amplified by PCR using the primers 5’-TTGAGGTAGTTGCTCCCAAAG-3’ and 5’-GGCATTCCTTGGTGGTCATA-3’, and the genotype was confirmed by DNA sequencing.

### Western blotting experiments

The Western blot analysis was performed as previously described (Sun et al., 2018). 500 *rnf2*^−/−^ and wild-type embryos were collected and dechorionated, and cell lysates were prepared using TNE buffer (50 mM Tris-HCl (pH 8.0), 150 mM NaCl, 1% Triton X-100, 0.5% sodium deoxycholate, 5 mM EDTA, 1× DTT) with the cOmplete proteinase inhibitor (Roche, Switzerland). The lysates were resolved on 15% SDS polyacrylamide gels and immunoblotted with primary antibody anti-Rnf2 (A302-869A, Bethyl, USA). The images were captured with the Image Quant LAS 4000mini (GE Healthcare Life Sciences, USA).

### Whole mount *in situ* hybridization (WISH)

WISH was performed as previously described (Sun et al., 2018). The DIG-labeled anti-sense probes were generated using DIG RNA Labeling Kit (SP6/T7) (Roche). The DNA templates used to generate probes were amplified by PCR and the primers used are listed in Table 1. For embryos at or after 36 hpf, the homozygotes were separated from their siblings according to their mobility and pectoral fin phenotype. For embryos before 36 hpf, as it was difficult to separate homozygous mutants from the heterozygous and wild type siblings, the WISH was performed for all progeny from the *rnf2+/−* parents. After the WISH, each embryo was photographed and genotyped. The photographs were taken under a stereomicroscope (Leica Z16 APO) with a digital camera (Leica DFC450, Germany).

**Table 1.**
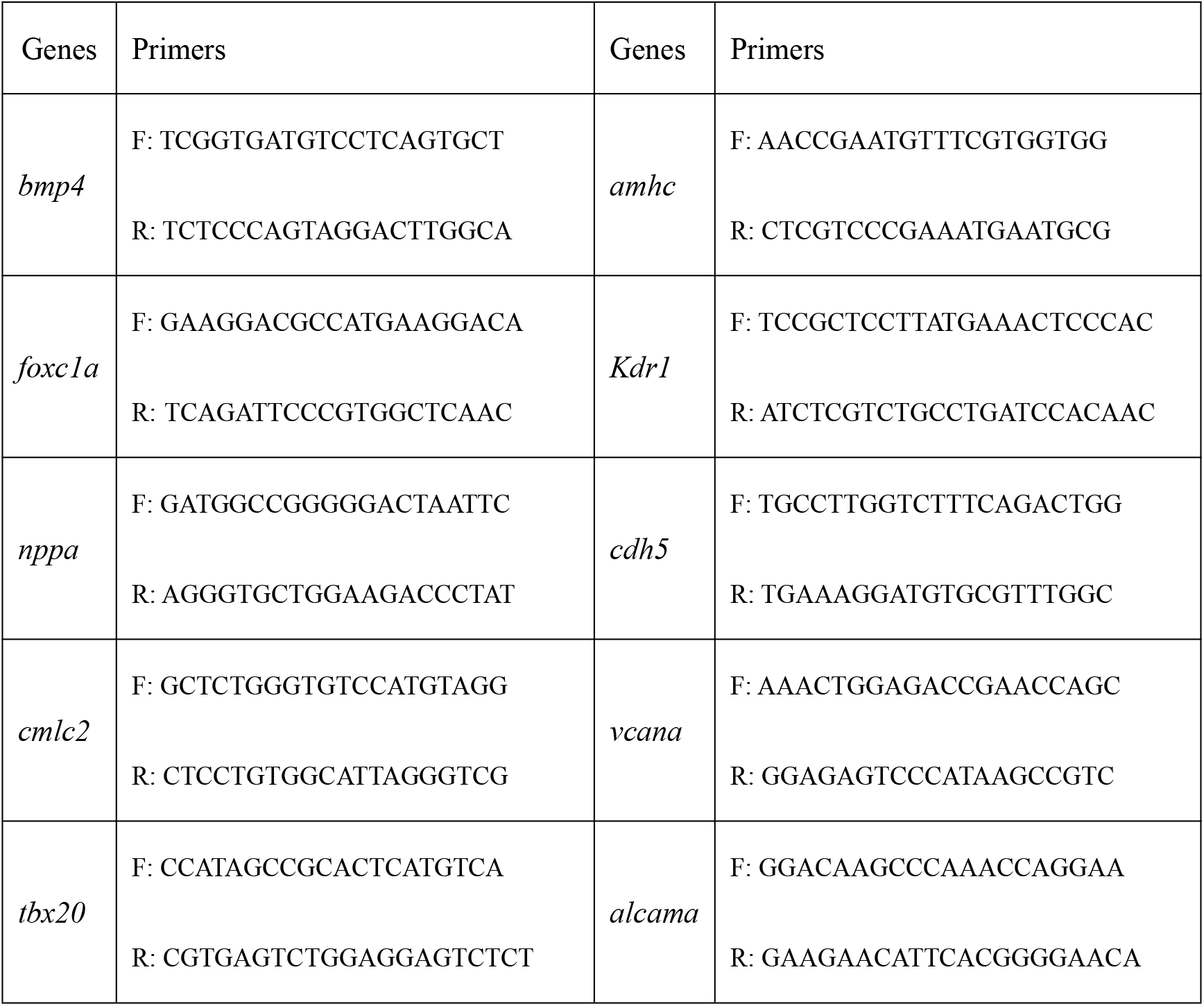
Primers used to amplify DNA template for Probe making

### Histological analysis

Zebrafish embryos at 76 hpf were fixed with Bouin’s solution at room temperature for 24h. The fixed embryos were embedded in 1% agarose, dehydrated with gradient ethanol, hyalinized with xylene and then embedded in paraffin. The paraffin embedded embryos were sectioned, and the sections were stained with the standard hematoxylin and eosin staining.

### Quantitative real time PCR

The relative mRNA levels of genes in *rnf2*^−/−^ and control hearts were detected by quantitative real time PCR (qRT-PCR). Primers used were listed in supplementary Table 2. Individual hearts from embryos were isolated for qRT-PCR analysis as described below. To collect the individual hearts, the transgenic zebrafish strain Tg (*myl7*: egfp) was used. The homozygous mutants were separated from their siblings according to their mobility and pectoral fin phenotype. The embryos were disrupted by pipetting up and down several times (depending on the developmental stages) with 10 µL pipette. Then the individual hearts were picked up by pipettes under a fluorescence stereomicroscope. Individual hearts were pooled into the same tube, and the total RNA was extracted using TransZol Up Plus RNA kit (Transgen, China). The cDNA was reverse-transcribed from the total RNA and was used for qRT-PCR analysis.

**Table 2.**
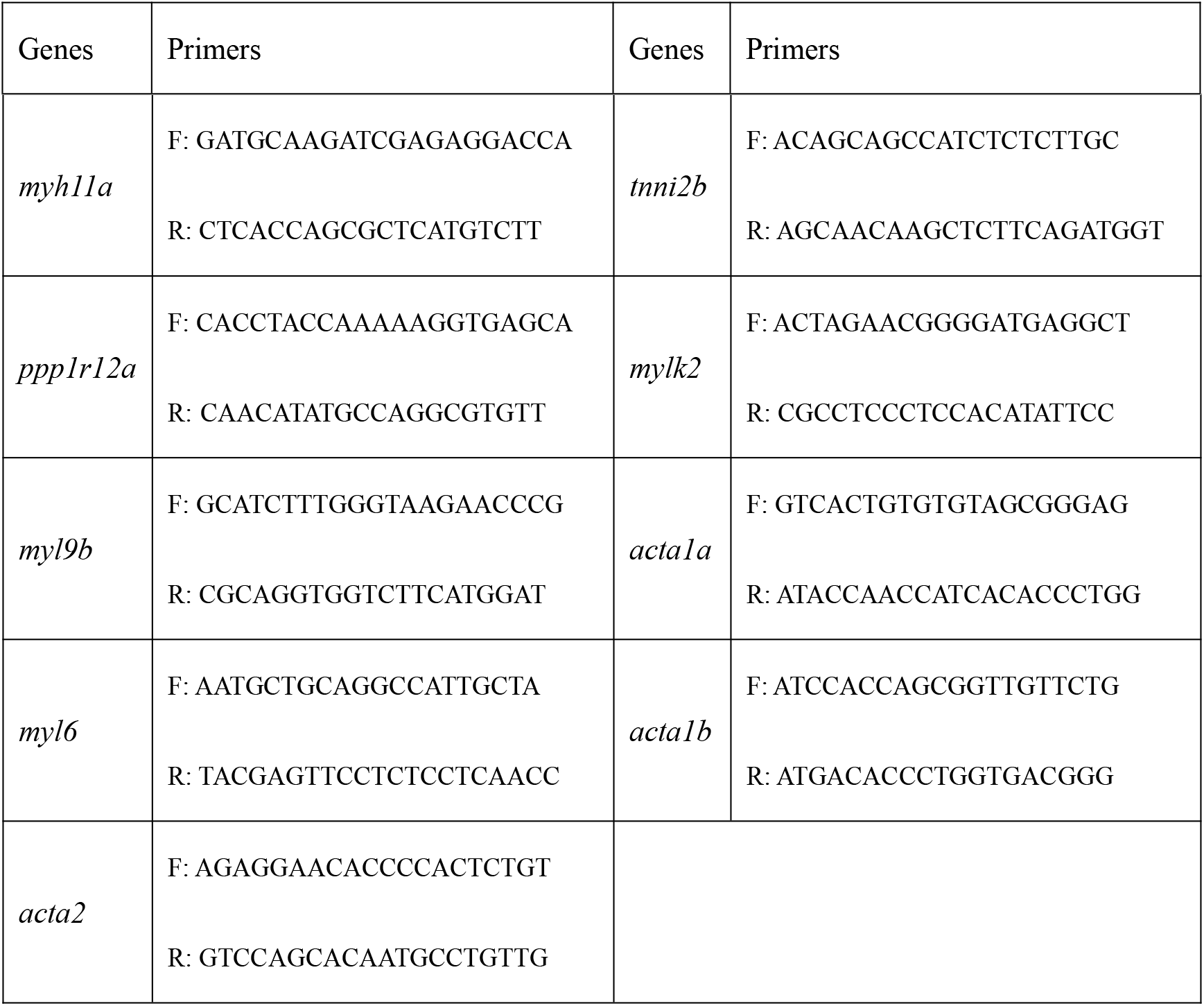
Primers for qRT-PCR

### Deep-RNA sequencing analysis of single hearts

The individual embryonic hearts were extracted as described above, and the remaining part of the embryo was collected separately and used for the genotyping. Single hearts from *rnf2*^−/−^ or wild-type embryos were collected and used for deep-RNA sequencing (RNA-seq). The sample library construction, RNA sequencing and preliminary data analysis were done by the Anoroad Gene Technology Company, China. An average of 40-50 M clean reads were obtained per heart, and the expression of approximately 18,000 genes was detected.

### ChIP-seq and ChIP-PCR experiments

The ChIP experiments were performed according to Agilent Mammalian ChIP-on-chip manual as described. Briefly, 500 embryos of 24 hpf embryos were de-chorionated and treated to single cells by grinding. The cells were fixed with 1% formaldehyde for 10 minutes at room temperature. Then the reactions were stopped by 0.125 M glycine for 5 min with rotating. The fixed chromatin were sonicated to an average of 200-500 bp (for ChIP-Seq) or 500-1000 bp (for ChIP-qPCR) using the Covaris Sonication System (USA) according to the manual. Then Triton X-100 was added to the sonicated chromatin solutions to a final concentration of 0.1%. After centrifuge, 50 µL of supernatants were saved as input. The remainder of the chromatin solution was incubated with Dynabeads previously coupled with 10 µg ChIP grade anti-Ring1b antibodies (A302-869A, Bethyl) overnight at 4℃ with rotation. Next day, after 7 times wash with the washing buffer, the complexes were reverse cross-linked overnight at 65℃. The ChIPed DNAs were extracted by hydroxybenzene-chloroform-Isoamyl alcohol and purified by a Phase Lock Gel (Tiangen, China).

For ChIP-PCR, the ChIPed DNAs were dissolved in 100 µL distilled water. Quantitative real-time PCR (qRT-PCR) was performed using a Bio-Rad qPCR instrument. The enrichment was calculated relative to the amount of input as described (Sun et al., 2020). All experiments were repeated at least three times. The relative gene expression levels were calculated based on the 2^−ΔΔCt^ method. Data were shown as means ± S.D. The Student’s t test was used for the statistical analysis. The significance is indicated as follows: *, *p < 0.05*; **, *p < 0.01*; ***, *p < 0.001*.

### Calcium signaling using Fluo 4-AM

The calcium signal was detected according to the kit manual. Briefly, the hearts isolated from 48 hpf *rnf2*^−/−^ and wild-type embryos were transferred to a 20 mm glass bottom cell culture dish containing external control solution ECS (140 mM NaCl, 4 mM KCl, 1.8 mM CaCl2, 1 mM MgCl2, 10 mM glucose, and 10 mM HEPES (pH 7.4). Then the embryos were incubated in ECS medium containing 5 µM Fluo 4-AM, 10 µM blebbistatin and 0.35 mM probenecid for 30 min. The blebbistatin was added to uncouple the excitation-contraction process in the zebrafish embryonic heart. After rinsed twice with ECS containing 0.35 mM probenecid, the hearts were transferred into ECS containing 0.35 mM probenecid. Images were taken with a confocal laser scanning microscopy (Leica SP8 DLS, Germany).

### Imaging, quantification and statistical analysis

The area of pericardial edema of 72 hpf embryos was measured by marking the edema region on photos using ImageJ. Heart rates were counted manually under the dissecting microscope (WPI). The heart beating frequency in one minute per embryo was calculated.

For transmission electron microscopy of cardiac muscle structure, 80 hpf *rnf2*^−/−^ and wild-type embryos were fixed, embedded in the Spur resin, and then sectioned. The sections were stained with uranyl acetate and lead citrate. Images were taken with a Hitachi TEM system (Japan). Data were analyzed with the GraphPad Prism 7.0 software. The values are presented as mean ± SEM. The *P* values were calculated by two-tailed Student’s test, * *p < 0.05*; ** *p < 0.01*; *** *p < 0.001*.

### Data availability

All data have been deposited in BIGD at the Beijing Genomic Institute (BGI): https://bigd.big.ac.cn/databases, under the accession numbers: CRA002288 (Rnf2 ChIP-seq data for 15- and 24-hpf embryos), CRA00176 (RNA-seq data for single hearts of 24 hpf embryos) and CRA002292 (RNA-seq data for single hearts of 36- and 48-hpf embryos).

## Results

### Generation of *rnf2−/−* mutants

The expression of *rnf2* during zebrafish embryogenesis has been described (van der Velden et al., 2012). *Rnf2* was maternally inherited, and ubiquitously expressed at pre-gastrulation, gastrulation and early somitogenesis stage. From 24 to 72 hpf, its expression was gradually confined to brain-, craniofacial-, pectoral fin-, gut- and heart regions, which implied that Rnf2 is persistently required for the development and homeostasis of these organs (Supplementary Fig. 1A-B).

To investigate the role of Rnf2 in zebrafish embryogenesis, we generated *rnf2* mutant lines using the CRISPR/Cas9 technology. The exon 3 of the *rnf2* gene was targeted and two mutant alleles *(rnf2^f5^* and *rnf2^f8^*) were generated as revealed by genotyping (Fig. 1A). The *rnf2^f5^* mutant allele carries a 5 bp (base pair) deletion in the exon 3 (at 115 bp) of *rnf2* gene. The *rnf2^f8^* mutant allele harbors an 8 bp deletion at the similar site. Rnf2 protein levels were not detected by Western blotting in 24 hpf *rnf2*^−/−^ embryos (Fig. 1B), suggesting that the mutant alleles were null. It is well known that Rnf2 functions as an E3 ubiquitin ligase that catalyzes H2AK119ubi (Van der Velden et al., 2012). Consistently, the levels of H2AK119ubi were greatly reduced in *rnf2*^−/−^ embryos compared to the sibling controls (Fig. 1B). We believed that the residual H2AK119ubi levels should be due to the maternal contribution of Rnf2. At 48 hpf mutant embryos, H2AK119ubi was hardly undetectable (data not shown; Van der Velden et al., 2012).

**Fig. 1.**
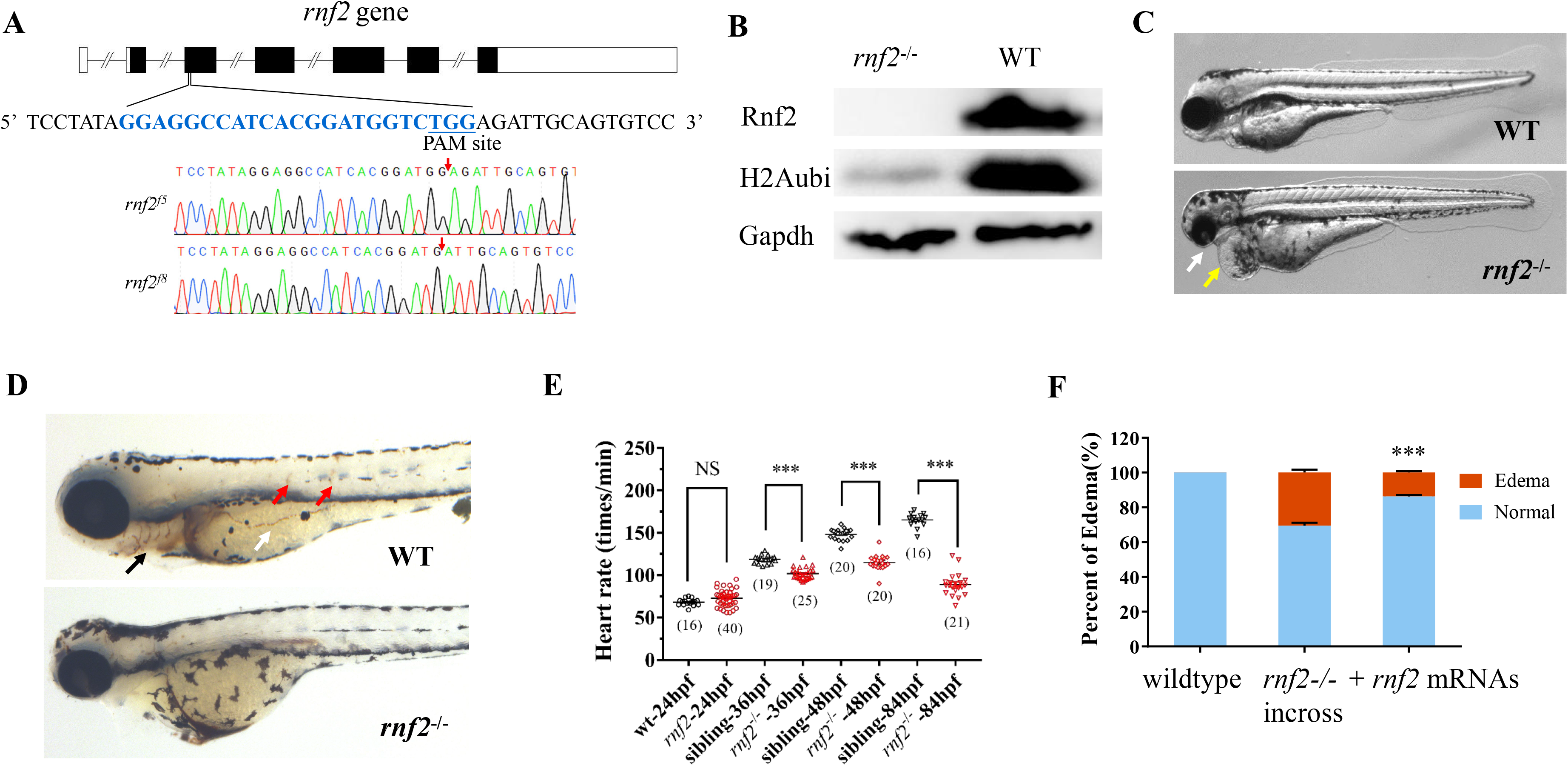
Generation of *rnf2* mutant zebrafish. **A** Cartoon showing the CRISPR/Cas9 targeting site and mutant alleles. **B** WB showing loss of Rnf2 and reduced H2Aub1 in 24 hpf *rnf2−/−* embryos. **C** Pleiotropic phenotypes of *rnf2−/−* mutant embryos at day 3, including small head, edema with a stringy heart, absence of pectoral fins and defective craniofacial structures. Yellow arrow showing edema, and white arrow showing the missing craniofacial structures. **D** o-Dianisidine staining for red blood cells in day 3 control and mutant embryos. Black arrows showing the blood signals in control embryos, which were missed or reduced in a portion of mutant embryos. **E** Quantification of heart rate of control and mutant embryos at the indicated time points. **F** Percentage of embryos showing edema or bradycardia for wildtype-, *rnf2+/−* incross and *rnf2+/−* incross embryos injected with *rnf2* mRNAs. *** indicates *p<* 0.001. All experiments were repeated three times and images are representatives.

There was no overt morphological differences between control and mutant embryos at 24 hpf, suggesting that *rnf2* is not strictly required for the formation of basic body plan in zebrafish (Chrispijn et al., 2019). However, *rnf2*^−/−^ embryos can be separated from their *rnf2*^+/+^ and *rnf2*^+/−^ siblings morphologically as early as 36 hpf, as they exhibited reduced pectoral fins with abnormal swimming manure. At 72 hpf, *rnf2*^−/−^ embryos usually exhibited pleiotropic phenotypes, including defective craniofacial structures, lack of pectoral fins and malformation of cardiovascular system (such as edema, bradycardia, hypotrophy and compromised blood flow) (Fig. 1C-D; Supplementary Fig. 1C-D). The roles of Rnf2 in craniofacial and pectoral fin development have been discussed (Van der Velden et al., 2012; 2013).

In this study, we focused on Rnf2 function in cardiovascular and blood development. Heart rate is a good readout for evaluating cardiovascular function. We carefully measured the heart rate of control and mutant embryos at 24-, 36-, 48- and 84 hpf. Although no overt difference was detected for 24 hpf embryos, a significant reduction in heart rate was readily detected at 36 hpf, and this became more marked at later time points, suggesting a progressing cardiovascular phenotype over time by loss of Rnf2 (Fig. 1E).

To verify that the cardiovascular defects were due to loss of Rnf2 function, rescue experiments were performed. Zebrafish *rnf2* or mouse *Rnf2* mRNAs were micro-injected into the 1-cell stage embryos from the heterozygous incross. Embryos with pericardial edema and bradycardia were counted and the edema area was measured at 72 hpf (Supplementary Fig. 1E). *Rnf2* mRNAs micro-injection reduced the percentage of embryos with edema and bradycardia from approximately 25% to 15% (Fig. 1F). Thus, our rescue experiments confirmed the specificity of *rnf2* mutant phenotypes.

### Abnormal up-regulation of hematoendothelial progenitor genes in *rnf2***^−/−^** embryos

Cardiovascular and hematopoietic progenitors are specified from mesodermal precursors during zebrafish gastrula stage (Paik and Zon, 2010). To investigate whether the observed cardiovascular defects were due to aberrant induction of early mesoderm, we collected 7 hpf embryos from *rnf2+/−* incross and performed WISH for a panel of mesodermal markers such as *eve1* (ventral mesoderm), *flh* (axial mesoderm) and *foxc1a* (paraxial mesoderm). The WISH results showed that the expression of these markers was largely comparable between the mutant and the control embryos (Supplementary Fig. 2A).

Shortly after gastrulation, cardiac, endothelial and blood precursors develop at the ALPM where early hematoendothelial progenitor genes such as *lmo2/tal1/etv2* and early cardiac specification genes such as *nkx2.5/hand2* are expressed in close proximity, with the former more anteriorly localized (Palencia-Desai et al., 2011). We therefore asked whether the expression of these genes was affected by loss of Rnf2 function, by performing WISH using embryos ranging from 3- to 18-somite stages. At 3- to 8-somite stages, the expression of hematoendothelial progenitor genes such as *lmo2/tal1/etv2* was slightly elevated, albeit in a small portion of the mutant embryos (Fig. 2A-B). As it is established that hematoendothelial progenitors and myocardial precursors antagonize each other at the ALPM, we performed two color WISH for *myoD*, *hand2* and *tal1* using 6-somite stage control and mutant embryos. The WISH results showed that the expression of *hand2* was slightly reduced while the expression of *tal1* was slightly expanded (Supplementary Fig. 2C). This observation was in line with the previous reports (Palencia-Desai et al., 2011; Schoenebeck et al., 2007). Between the 15- and 18-somite stages, the *tal1/lmo2*/*etv2* expressing hematoendothelial/endocardial precursors migrate medially and posteriorly, and arrive at the midline. When examining 18-somite stage embryos, we found that *tal1/lmo2/etv2* expression levels remained slightly higher in the mutant embryos than in the control embryos (Fig. 2C).

**Fig. 2.**
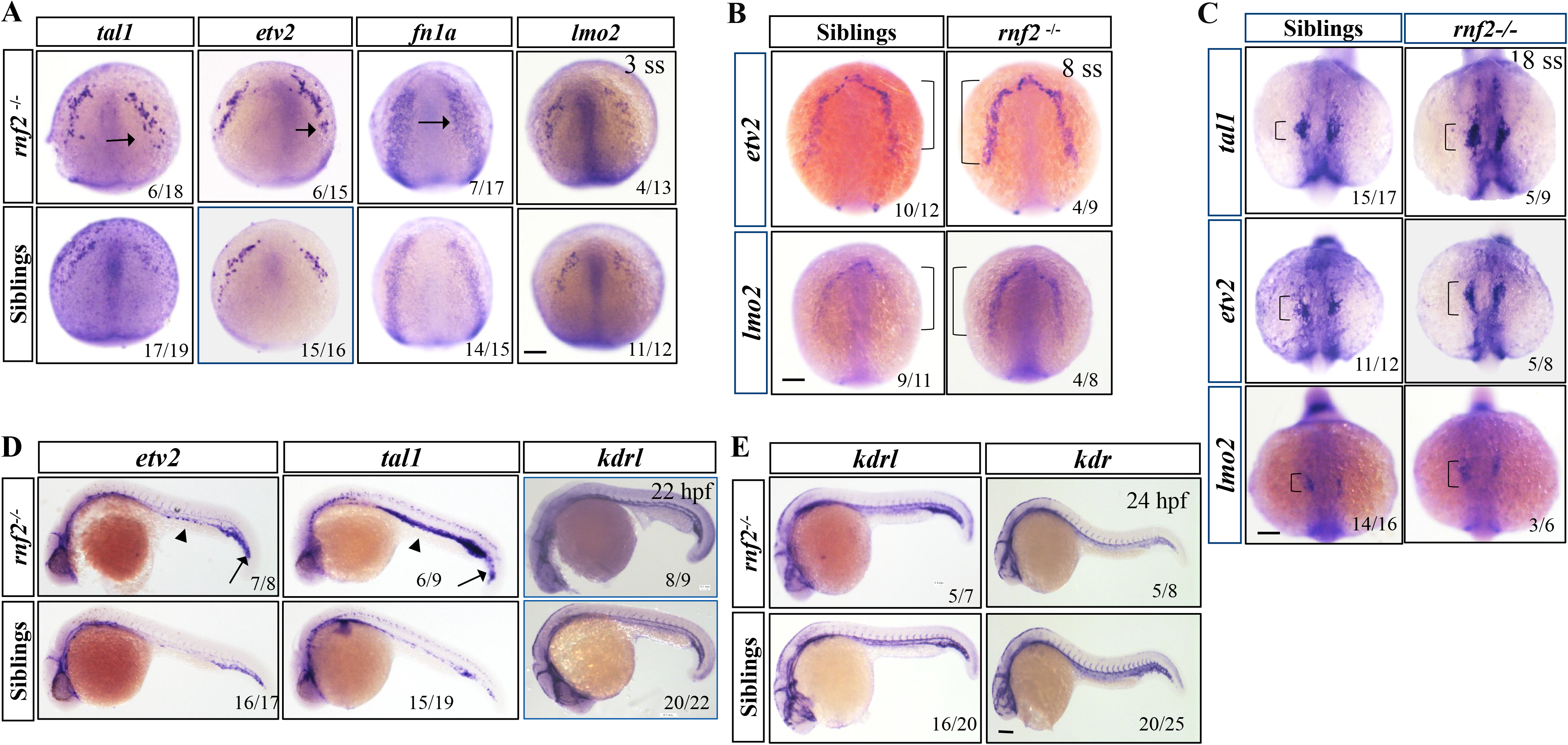
Abnormal up-regulation of hematoendothelial genes in mutant embryos. **A** WISH results for control and mutant embryos showing the expression of hematoendothelial genes *tal1*, *etv2, fn1a* and *lmo2* at 3 somite stage. Scale bar: 0.2 mm. **B** WISH results showing the expression of *etv2* and *lmo2* in 8 somite stage embryos. Brackets were used to indicate the expression domains of the genes. Scale bar: 0.2 mm. **C** WISH results showing the expression of *etv2*, *tal1* and *lmo2* in 18 somite stage embryos. Brackets were used to indicate the expression domains of the markers. Scale bar: 0.2 mm. **D** WISH results showing the expression of *etv2*, *tal1* and *kdrl* in control and mutant embryos at 22 hpf. Arrow heads showing the enhanced expression of *etv2* and *tal1* in the intermediate cell mass in mutants compared to the controls. **E** WISH results showing the expression of *kdr* and *kdrl* in control and mutant embryos at 24 hpf. Scale bar: 0.2 mm. All experiments were repeated three times and images are representatives.

Next, we went on to examine the expression of hematoendothelial progenitor and myocardial genes for control and mutant embryos at 22-30 hpf. *Etv2* and *tal1* expression levels in a variety of hematoendothelial derivatives, including the cranial vasculature, the intermediate cell mass (ICM), the posterior cardinal vein and the tail plexus region, were clearly higher in mutant embryos than in controls (Fig. 2D). It is well-known that the primitive wave of hematopoiesis originates from the ICM. This observation suggested that generation of *etv2*-expressing primitive HSCs may be enhanced in the absence of Rnf2. As expected, the expression of cardiac chamber genes such as *nkx2.5*, *nppa/b*, *amhc*, *vmhc* and *tbx20* was slightly reduced (Supplementary Fig. 2D-E) (Schupp et la., 2014). The formation of blood vessels appeared grossly normal in the absence of Rnf2, based on the expression of vascular endothelial markers such as *kdr*, *kdrl* and *cdh5* (Fig. 2D-E; Supplementary Fig.2F).

Taken together, we concluded that the hematoendothelial progenitor genes were up-regulated and the cardiac chamber genes were slightly down-regulated in *rnf2* mutant embryos, compared to the control embryos.

### Primitive wave of hematopoiesis was compromised in *rnf2* mutant embryos

In parallel with cardiovasculogenesis, primitive hematopoiesis is initiated from a specialized subset of developing vascular endothelium (called hemogenic endothelium) and can give rise to HSCs (Paik and Zon, 2010). It is well-known that transcription factors such as Tal1/Lmo2 and Etv2 function at the top of the regulatory cascade that is essential for hematopoietic development (Gering et al., 2003; Liu et al., 2019; Sumanas and Lin, 2006), and that PcG component proteins repress gene expression for proper HSC regulation and function (Di Carlo et al., 2018). The abnormal expression of master hematopoietic TF genes such as *tal1/lmo2/etv2* prompted us to investigate whether there were HSC-related defects in *rnf2* mutant embryos.

First, we investigated the primitive wave of hematopoiesis which gives rise to primitive erythrocytes, macrophages and neutrophils in zebrafish. *Gata1*/*hbbe1*, *pu.1/mpeg1* and *mpx/lyz* are well-known early erythroid-, myeloid- and granulocyte- markers, respectively (Lian et al., 2018; Liu et al., 2019). WISH experiments were performed for *gata1/hbbe1*, *pu.1/mpeg1*, and *mpx/lyz*, using mutant and control embryos at 24 and 48 hpf. The WISH results showed that the expression of early erythroid marker *gata1* was increased in mutant embryos compared to controls, while the expression of myeloid- and granulocyte-markers was obviously decreased (Fig. 3A-B; Supplementary Fig. 3A-B).

**Fig. 3.**
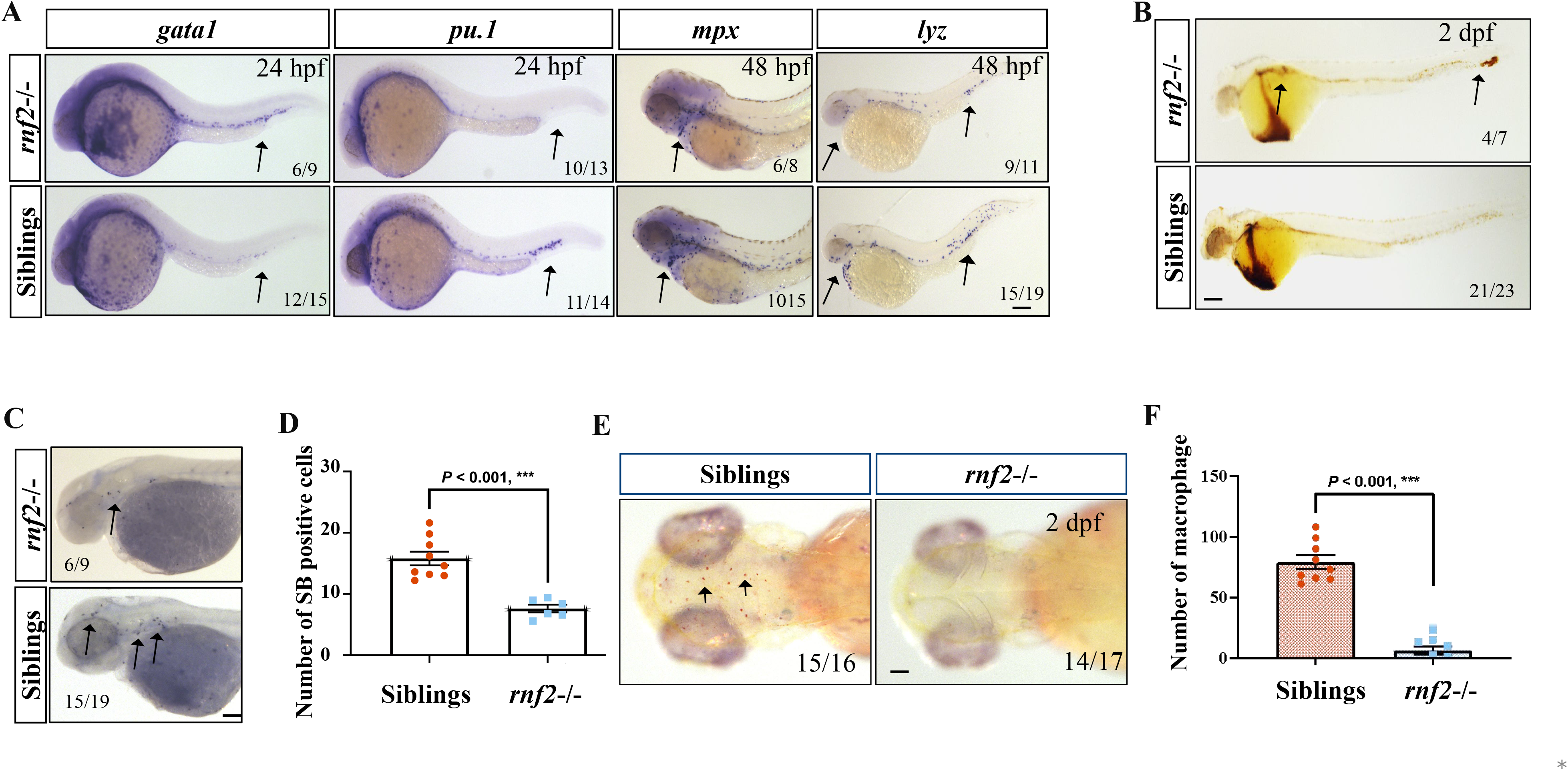
Primitive wave of hematopoiesis was defective in rnf2−/− embryos. **A** WISH results of control and mutant embryos, showing the expression of hematopoietic markers *gata1*, *pu.1, lyz* and *mpx*. *Mpx* signals were shown at head and yolk sac regions. Scale bar: 0.2 mm. **B** o-Dianisidine staining of red blood cells in 2 dpf control and mutant embryos. Arrows showing the slightly increased o-Dianisidine positive cells in mutant embryos compared to the controls. **C** Sudan black B (SB) staining of neutrophils for 2 dpf control and mutant embryos. Arrows showing the SB positive signals at head and yolk sac regions. **D** Quantification for panel C. **E** NR staining of macrophage for 2 dpf control and mutant embryos. Arrows showing the NR staining positive cells at head regions. **F** Quantification for panel E. All experiments were repeated three times and images are representatives.

The WISH results strongly suggested that initiation of primitive hematopoiesis was affected in *rnf2* mutant embryos. To substantiate this, we performed NR staining (inflammatory macrophage migration) for macrophage, o-Dianisidine staining for red blood cells and Sudan black B (SB) staining for neutrophils (Lian et al., 2018). o-Dianisidine staining results confirmed that the erythrocyte numbers were slightly elevated (Fig. 3B). As shown in Figure 3C-F, the number of macrophage and neutrophils was greatly reduced in the mutant embryos compared to the control embryos. In particular, differentiation into macrophage was almost completely blocked in the absence of Rnf2 (Fig. 3E-F).

### Definitive wave of hematopoiesis was defective in *rnf2* mutant embryos

Next, we asked whether the definitive hematopoiesis was affected in *rnf2* mutants. To this end, we first determined the expression of *c-myb* and *runx1* that mark the definitive HSCs using day 2 control and mutant embryos. In zebrafish, HSCs colonize the caudal hematopoietic tissue (CHT) transiently before seeding the definitive hematopoietic organs (Paik and Zon, 2010). The WISH results showed that the *rnf2* mutant embryos displayed more *c-myb* and *runx1* positive cells in the CHT, compared to the control embryos (Fig. 4A). This observation suggested that generation of definitive HSCs was enhanced in the absence of Rnf2.

**Fig. 4.**
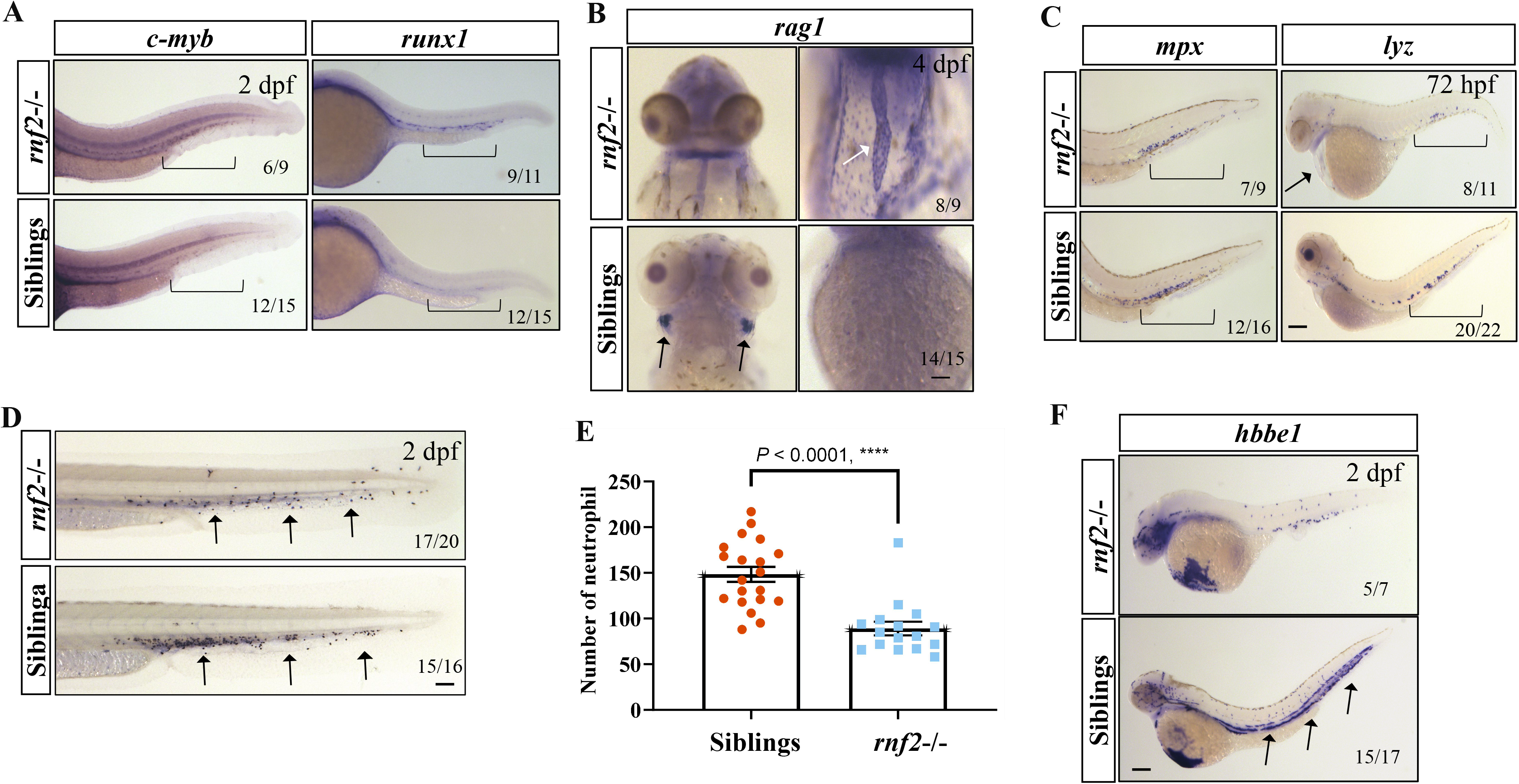
Definitive wave of hematopoiesis was defective in rnf2−/− embryos. **A** WISH results for *c-myb* and *runx1*, showing slight increased both *c-myb* and *runx1* positive HSCs in CHT of mutant embryos compared to the controls. Brackets indicate the CHT region. **B** WISH results showing the expression of *rag1*. Note that *rag1* is strongly expressed in the thymus of control but not mutant embryos (left, dorsal view), and is ectopically expressed in the mutant hearts but not in control hearts (right). Black arrows showing *rag1* expressing thymus in the control embryo, and white arrows showing *rag1* expressing hearts in the mutant embryo. **C** WISH results for *mpx* and *lyz* of control and mutant embryos. Brackets showing the CHT region. **D** SB staining for neutrophils. Black arrows showing the SB positive cells. **E** Quantification for D. **F** WISH results for the erythrocyte marker *hbbe1.* Black arrows showing the *hbbe1* positive blood cells. All experiments were repeated three times and images are representatives.

The definitive wave of hematopoiesis generates all the different types of circulating blood cells, including erythrocytes, myeloid cells (macrophage and neutrophil) and lymphocytes. To understand the role of Rnf2 in definitive hematopoiesis, we first investigated the lymphoid lineage by performing WISH for *rag1*, a definitive lymphoid marker. Compared to the control embryos, *rnf2* mutants at 4 dpf almost lacked of *rag1* positive signals in the nascent thymus (Fig. 4B; Supplementary Fig. 4A). Surprisingly, we found that *rag1* was ectopicaly expressed in the mutant hearts, whereas it was never expressed in the control hearts.

Next, WISH experiments were performed for *mpx* as well as *lyz*, a definitive neutrophil marker. The results showed that the expression of both markers was reduced in mutant embryos compared to the controls (Fig. 4C; Supplementary Fig. 4A). To confirm this, we determined the neutrophil numbers by the SB staining (Fig. 4D). Quantification of the positive cells in the trunk, tail and CHT showed that the number of neutrophils was indeed greatly reduced (nearly half of the controls) (Fig. 4E).

To assess erythropoiesis, we performed WISH for *hbbe1* using day 2 control and mutant embryos. The results showed that generation of red blood cells was severely compromised in the absence of Rnf2 (Fig. 4F). o-Dianisidine staining for red blood cells gave a similar result (Supplementary Fig. 4B).

Taken together, we concluded that Rnf2 is absolutely required for normal embryonic hematopoiesis, and its loss leads to severe defects in both primitive and definitive waves of hematopoiesis during zebrafish embryogenesis.

### Rnf2 is persistently required to stabilize cardiac gene expression by repressing hematoendothelial progenitor genes

*Rnf2* is clearly expressed in the heart of 48-72 hpf embryos (Supplementary Fig. 1B), suggesting that sustained activities of Rnf2 are required for cardiac maturation and homeostasis. We speculated that Rnf2-mediated epigenetic mechanism may continue to repress the hematoendothelial progenitor genes in the differentiated heart.

To investigate this, we performed WISH using an expanded panel of hematoendothelial progenitor markers, including *gata2*, *tal1*, *lmo2*, *nfatc1*, *etv2, fn1*, *fev*, *hoxa3* and *hoxb4*, for 48 hpf control and mutant embryos (Fig. 5A; Supplementary Fig.5A). Globally, all these marker genes appeared ectopically overexpressed in mutant hearts compared to the control counterparts. Specifically, the expression domain of *nfatc1* gene was primarily restricted around the AVC region in the control hearts, while it was expanded to the entire endocardium in the mutant hearts. *Etv2* expression at the time was primarily confined to the ventricular part of endocardium in the control hearts, while it was distributed throughout the entire endocardium in the mutant hearts. Of note, hematopoietic-specific TF genes such as *gata2/hoxa3/hoxb4a* were barely detected in 48 hpf control hearts. By contrast, they were readily expressed in the mutant hearts. Thus, Rnf2 is persistently required to repress the master hematoendothelial TF genes, which is likely important for cardiac maturation and homeostasis.

**Fig. 5.**
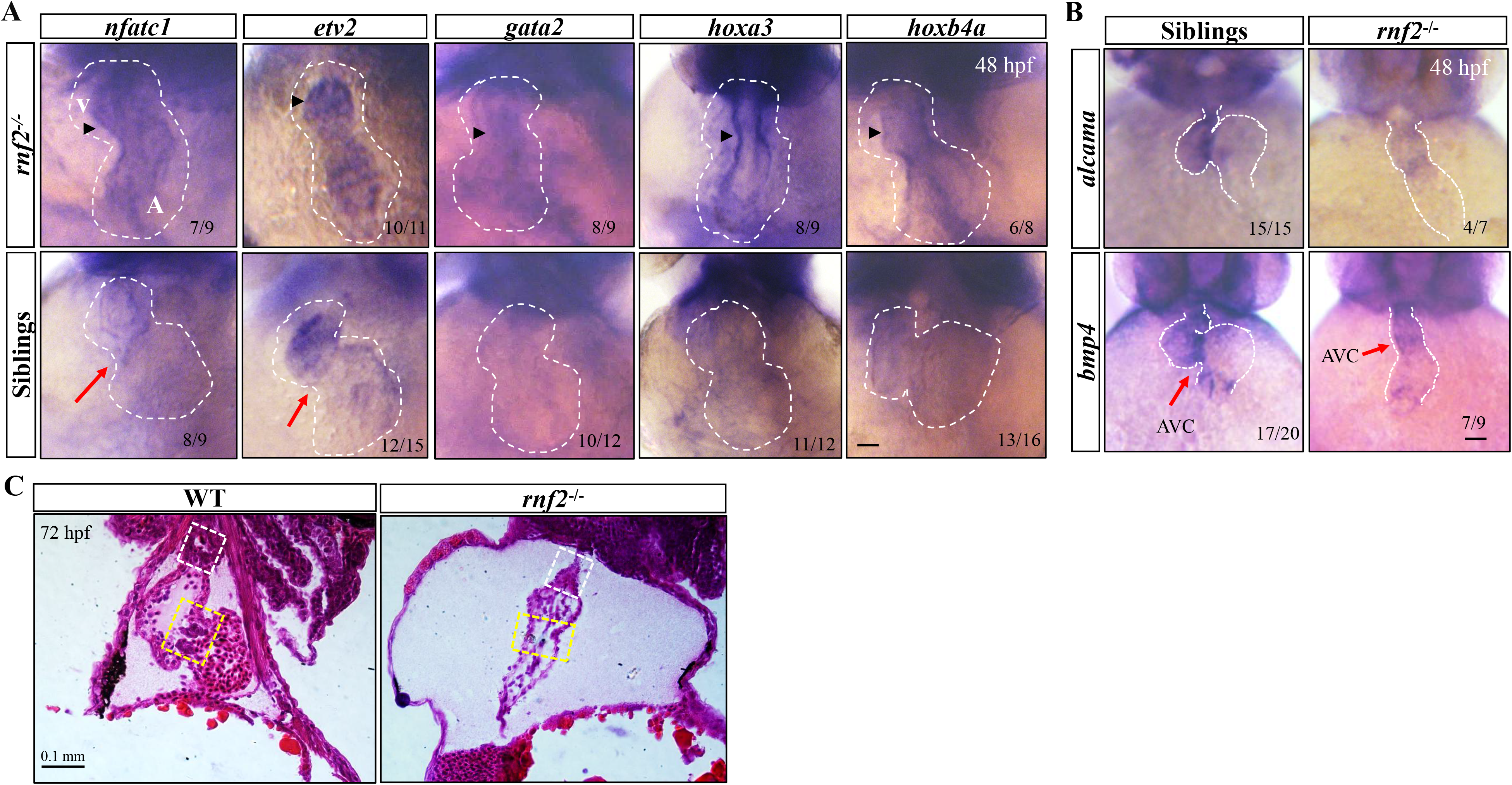
Ectopic expression of hematoendothelial markers in mutant hearts. **A** WISH results of control and mutant embryos, showing the expression of the indicated markers. Black arrow head indicates the endocardium, and red arrow showing the AVC region between the atrial and the ventricular. The white dashed line showing the morphology of the hearts. Note the defective cardiac looping in mutant embryos. V: ventricular; A: atrial. Scale bar: 0.1 mm. **B** WISH results showing the reduced expression of AVC markers in mutant hearts. The white dashed line showing the morphology of the hearts. Note the defective cardiac looping in mutant embryos. Red arrows pointing to the AVC region. Scale bar: 0.1 mm. **C** HE staining of hearts from *rnf2−/−* and wild-type embryos. White dotted boxes indicate the bulbus arteriosus, and yellow dotted boxes indicate the AVC and primitive valve regions. Note the defective formation of both bulbus arteriosus and primitive valve in the mutant heart. All experiments were repeated three times and images are representatives.

Heart valves originate from a subpopulation of endocardium called endocardial cushion cells (ECCs) located at defined regions of the atrial-ventricular canal (AVC) and the outflow tract (OFT) (Puceat, 2013). Combing the abnormal expression of hematoendothelial TF genes in endocardium, the cardiac looping defect as well as the bradycardia phenotype in *rnf2* mutant embryos, we speculated that AVC and primitive valve formation might be defective by loss of Rnf2 function. The specification of AVC can be marked by the restricted expression of *vcana* and *has* in endocardium as well as of *bmp4* and *alcama* in myocardium. The endocardial cushion and AV canal marker *has2* was normally expressed at 1-2 somite stage (Supplementary Fig. 5B). We examined the expression of *vcana*, *bmp4* and *alcama* by performing WISH using control and mutant embryos at 48 hpf (Fig. 5B; Supplementary Fig. 5C). In control embryos, the AVC was well-formed between the atrial and ventricular chambers, and the expression of *bmp4, alcama* and *vcana* was restricted to the AVC region. By contrast, *rnf2*^−/−^ embryos lacked of a clear AVC structure, and the expression of *bmp4* and *alcama* was sparse and diffuse, displaying a loss of AVC-restricted expression pattern. Although *vcana* remained at the AVC region, its expression levels were much reduced in mutant hearts. These observations suggested that the AVC specification was indeed affected in the absence of Rnf2.

To investigate whether the primitive valve formation was defective in *rnf2−/−* embryos, sections of hearts from 72 hpf control and mutant embryos were subjected to the HE staining. Globally, the heart was smaller and hypotrophy in mutant embryos compared to the controls. In control embryos, the protruding primitive valve leaflets were well-formed, comprising of morphologicaly distinct layer of valvular edothelial cells and interstitial cells (Fig. 5C). By contrast, the primitive valve leaflets in mutant embryos was severely reduced or missing, exhibiting few valvular interstitial cells.

Taken together, we concluded that Rnf2 is persistently required for the stabilization of cardiac gene program, cardiac maturation and the proper formation of AVC and primitive valve.

### Transcriptome comparison assay for control and *rnf2* mutant embryos

The above data showed that a few of hematoendothelial progenitor genes were deregulated in *rnf2* mutant embryos, which may lead to defective hematopoiesis and cardiogenesis in zebrafish embryos. To comprehensively understand the underlying mechanism by which Rnf2 regulates cardiovascular and hematopoietic development, deep RNA-sequencing was performed for control and *rnf2−/−* embryos at 24 hpf, as well as for individual hearts isolated from control and *rnf2−/−* embryos at 24 and 48 hpf (Fig. 6A).

**Fig. 6.**
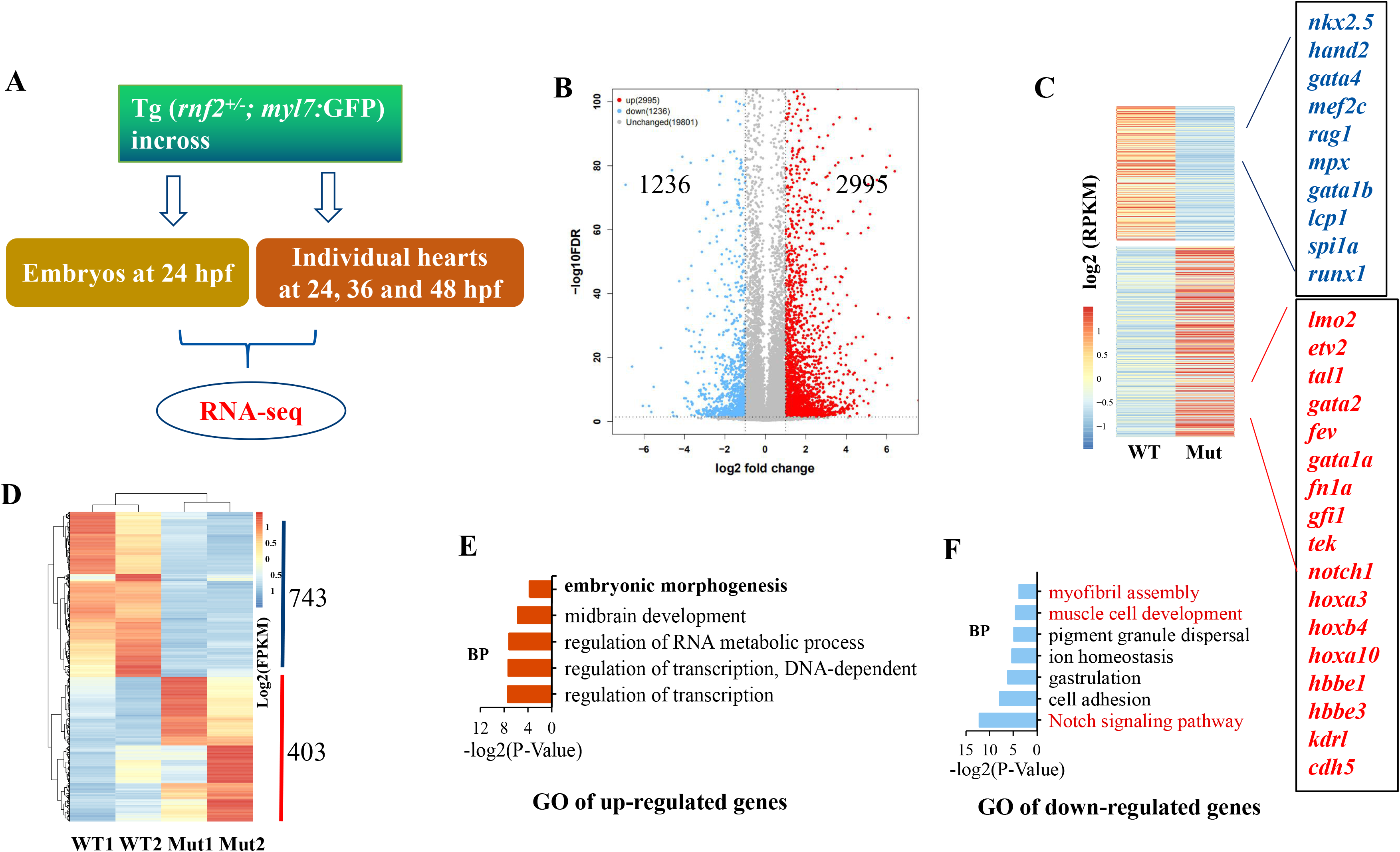
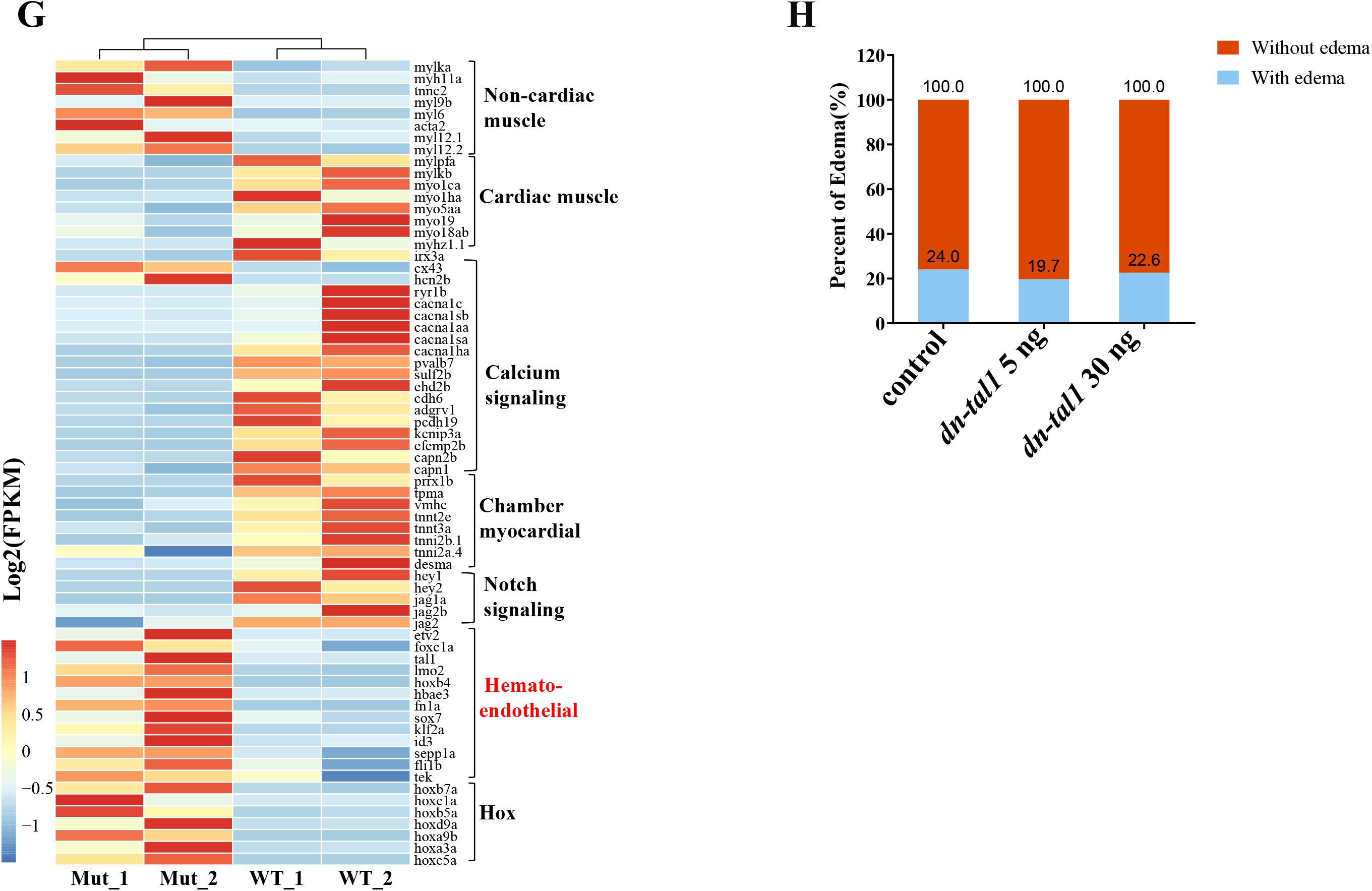
Transcriptome analysis of single hearts isolated from control and mutant embryos. **A** Cartoon showing the deign of the RNA-seq experiments. **B** valcano plot showing the up- and down-regulated genes for 24 control and mutant embryos. Fold change of Log2 (*rnf2−/−*/control). **C** Heat map showing representative DEGs in B. Boxes at right showing the representative cardiac- and hematoendothelial-genes that were up- or down-regulated in *rnf2* mutant embryos. WT: for wildtype embryos; Mut: for *rnf2* mutant embryos. **D** Heat map showing the up- and down-regulated DEGs from individual hearts of control and mutant embryos at 24 hpf. WT1: replicate 1 for wildtype embryos; Mut1: replicate 1 for *rnf2* mutant embryos. **E-F** GO analysis of up-regulated (C) and down-regulated (D) genes in 24 h *rnf2−/−* hearts. **G** Heat map showing the up- and down-regulated genes, highlighting non-cardiac muscle-, cardiac muscle-, calcium signaling-, chamber myocardial-, hematoendothelial- and *hox* family genes. **H** Percentage of edema of in *rnf2+/−* incross that were injected with or without different dose of *dn-tal1* mRNAs. NS: no significant; ***, *P≤* 0.001.

First, we analyzed the RNA-seq data for control and mutant embryos at 24 hpf. A total of 4,231 differentially expressed genes (DEGs) were identified (log2 fold change> 1; *P*< 0.05). Of which, 2,995 genes were up-regulated and 1,236 genes were down-regulated (Fig. 6B). KEGG analysis showed that the up-regulated DEGs were enrichment for terms such as ECM-receptor interaction, focal adhesion, cardiac muscle contraction and dilated cardiomyopathy, and the down-regulated DEGs for terms such as regulation of lipolysis and PPAR signaling pathways (Supplementary Fig. 6A-B). As PcGs are known to primarily function as transcriptional repressors, we focused on the up-regulated DEGs (Ku et al., 2008; Rougeot et al., 2019). Hox family of transcription factor genes, which are the putative PRC1/Rnf2 targets, were markedly up-regulated (Blackledge et al., 2014). Hbbe family genes hbbe1/3 as well as hematoendothelial-lineage genes such as *etv2*, *tal1*, *gata2*, *tek*, *fe*v, *lmo2*, *gata1a*, *hoxb4,* gfi1, *fn1a* and *notch1*, were significantly up-regulated. By contrast, early cardiac specification genes such as *isl1*, *nkx2.5/2.7, mef2c*, *hand2*, *gata4/5/6* and *tbx5/20*, as well as hematopoietic genes such as *rag1, mpx, gata1b, lcp1(l-plastin), spi1a and runx1* was down-regulated. A heat map for the DEGs was shown, with representative up- and down-regulated genes listed aside in blue and red, respectively (Fig. 6C).

Next, RNA-seq data were analyzed for hearts isolated from control and mutant embryos. A total of 1,146 differentially expressed genes (DEGs) were identified from hearts of control and mutant embryos at 24 hpf. Of which, 403 genes were up-regulated and 743 genes were down-regulated (log2 fold change> 1; *P*< 0.05) (Fig. 6D). Approximately 1,124 DEGs were identified from hearts of 48 hpf embryos. Of which, 712 genes were up-regulated and 412 genes were down-regulated (log2 fold change> 1; *P*< 0.05). A representative heat map of all DEGs for four individual hearts (from two mutant and two control embryos at 24 hpf) was shown, showing the reproducibility of our RNA-seq experiments (Fig. 6D). Gene ontology (GO) analysis revealed that the up-regulated DEGs were enriched for terms such as neural and brain development, DNA-binding and regulation of transcription (Fig. 6E), and the top terms for down-regulated DEGs were myofibril assembly, cardiac muscle development, ion homeostasis and the Notch signaling pathway (Fig 6F). Specifically, the endocardial/hematoendothelial progenitor TF genes, including *etv2*, *nfatc1*, *tal1* and *lmo2*, were significantly up-regulated at both time points (fold change> 2; *P*< 0.05), which coincides with our WISH experiments (Fig. 5A). In addition, we found that the expression of hematopoietic-specific genes such as *gata2*, *hhex*, *hoxa3*, *hoxb4*, *rag1*, *fev*, *pu.1/ spi1a*, *mpx, runx1*, *lcp1* and *mpeg1* was also markedly elevated. By contrast, the expression of early cardiac specification genes such as *Isl1*, *nkx2.5/2.7*, *hand2*, *gata4/5/6* and *tbx5/20* was barely altered (fold change< 1.5, and show no significant in statistics), which was in line with the previous report (Chrispijn et al., 2019). The qRT-PCR analysis of selected genes confirmed the RNA-seq results (Supplementary Fig. 6C-E).

If the master regulators of hematopoietic and vascular endothelial genes were up-regulated in the mutant hearts, the expression of their downstream genes should follow the same trend. In fact, we found that this was the case, based on the the RNA-seq data. Hematoendothelial related genes such as *alas2*, *crip2*, *dab2*, *epor*, *edn1/2*, *egfl7*, *eng1*, *erg*, *esam*, *fev, fli1*, *foxc1a*, *fn1a*, *gfi1*, *hoxa9*, *id3*, *klf2a*, *mef2ca*, *mrc1a*, *selp*, *sox7*, *sox17*, *tie1*, *tek, vegf*, *vwf*, *zfpm2b* and *cxcl12/cxcr4* were all up-regulated to a different extent (fold change> 1.4). On the other hand, *hey2* and *flt1*, which are known negatively-regulated by Tal1, were significantly down-regulated (Bussmann et al., 2007).

We also analyzed the down-regulated DEGs identified from hearts of control and mutant embryos. Loss of Rnf2 led to the down-regulation of many sarcomeric- and cardiac muscle-specific genes, including cardiac *troponin T* (*tnnt1*, *tnnt2e* and *tnnt3a*) and *I* (*tnni2* and *tnni1*), *ttn*, *nppa, nppb, actn1-3*, *prrx1b*, *lmna*, *tpma*, *desmin, smyhc1*, *vcan, vmhc*, *six1a*, *myh7*, *mylpfa*, *myo1*, *myo5*, *myo18/19*, *mylk5*, *mylkb*, *myhz1* and *myh14* (fold change> 1.5; *P*< 0.05) (Fig. 6G). Of which *tnnts*, *lmna, ttn, myh7*, *desmin* and *tpma* are known to be associated with dilated cardiomyopathy or bradycardia. Genes involved in calcium handling such as *ryr2b*, *cav1*, *cacna1g*, *cacna1c*, *cacna1s*, *cacna1b, cacna1ha*, *calsequestrin* (*casq1b*) and *parvalbumin* (*pvalb7*) were significantly down-regulated (fold change> 2; *P*< 0.05). Furthermore, key cardiac conduction regulatory genes such as *id2* and *irx3* were also down-regulated, accompanying with the deregulation of conduction-related genes such as *tbx2/3*, *cx43*, *hcn2* and *hcn4* (log2 fold change> 1; *P*< 0.05) (van Weerd and Christoffels, 2016). Notch pathway components such as *jag1/2* as well as its target genes *hey1/2* and *her1/2/4*, were significantly down-regulated. Notch signaling pathway is known a critical regulator for hematopoietic stem and progenitor cell specification (Liu et al., 2019). In addition, the profibrotic genes such as *postn* and *tgfb3* were down-regulated, which may contribute to the cardiac hypotrophy phenotype of the mutant embryos (Delgado-Olguin et al., 2012). Thus, loss of Rnf2 leads to down-regulation of genes involved in myofibril assembly, cardiac muscle development, cardiac conduction system, calcium handling and the Notch signaling.

As overexpression of hematoendothelial progenitor genes have been shown to inhibit myocardial and muscle cell fates in both in vitro and in vivo models (Castano et al., 2019; Veldman et al., 2012; Palencia-Desai et al., 2011; Patterson et al., 2007), we speculated that the reduced expression of chamber genes and compromised cardiac function might be at least partially due to the up-regulation of hematoendothelial progenitor genes. If this was the case, restoring normal activities of key hematoendothelial genes should rescue the gene expression and functional defects of *rnf2* mutant hearts. To investigate this, we reduced *tal1/gata2* gene activities by injecting dominant-negative form of *gata2* or *tal1* mRNAs (*dn-gata2* or *dn-tal1*) into the 1-cell stage embryos from *rnf2* heterozygous incross. The dominant-negative form of *Gata2* or *Tal1* has been previously described (Dasen et al., 1999; Lazrak et al., 2003). The results showed that injection of lower dose of *dn-tal1* or *dn-gata2* mRNAs (5-10 ng) can partially rescue the edema phenotype (Fig. 6H). Injection of higher dose of *dn-tal1* (30 ng) had a minimal rescue effect, suggesting that *tal1* activities must be in a narrow range. Injection of higher dose of *dn-gata2* mRNAs (30 ng), however, caused severe dorsal-ventral patterning defects of 24 hpf embryos, which prevented accurate evaluation of rescue effects on cardiac phenotypes.

Combining the RNA-seq data from whole embryos and single hearts as well as the WISH results, we concluded that Rnf2 is required for suppressing key hematoendothelial progenitor genes in cardiovascular precursors in the ALPM and the differentiated descendants (including the heart and HSCs), and its loss lead to abnormal up-regulation of these genes, resulting in disruption of normal cardiovascular and hematopoietic development.

### Hematoendothelial progenitor genes are directly regulated by Rnf2

To understand the molecular mechanism by which Rnf2 controls gene expression, we performed ChIP-seq experiments for 15 and 24 hpf embryos using Rnf2 antibodies. An average of 2,027 binding sites were identified genome-wide, based on two Rnf2 ChIP-seq replicates. Rnf2 peaks were mostly distributed at gene promoter and gene body regions (Fig. 7A-B). Bioinformatics analysis revealed that approximately 500 genes were highly enriched for Rnf2 signals, most of which encode developmental regulators, including Hox (homeodomain), Tbx (T-box), Nkx, Gata, Irx, Pax, Dlx and Sox families of transcription factors (Table 3). It is known that developmental regulator genes are in a poised state in stem cells, as they are occupied by both active mark H3K4me3 and repressive H3K27me3/H2Aub1 marks in ESCs (Ku et al., 2008). Interestingly, H2Aub1-positive bivalent promoters more efficiently retain H3K27me3 upon differentiation than PRC1-negative counterparts, which suggested that PRC1/RNF2 play important roles in the control of gene expression in conjunction with PRC2/EZH2. We found that developmental regulatory genes, including early cardiac specification genes such as *nkx2.5/2.7*, *gata4/5/6*, *isl1* and *tbx5* as well as hematoendothelial progenitor genes such as *gata2*, *hoxa3/b4*, *tal1 lmo2* and *etv2,* displayed broad Rnf2 ChIP-seq signals at promoter regions (Fig. 7C; Supplementary Fig. 7A), which implied that Rnf2 is important to repress their expression in zebrafish. Gene ontology (GO) analysis of Rnf2-bound genes showed that the top molecular function was related to DNA-binding and the regulation of transcription, and that the top biological processes were related to embryonic brain development, eye development, neuronal differentiation and heart development (Fig. 7D).

**Fig. 7.**
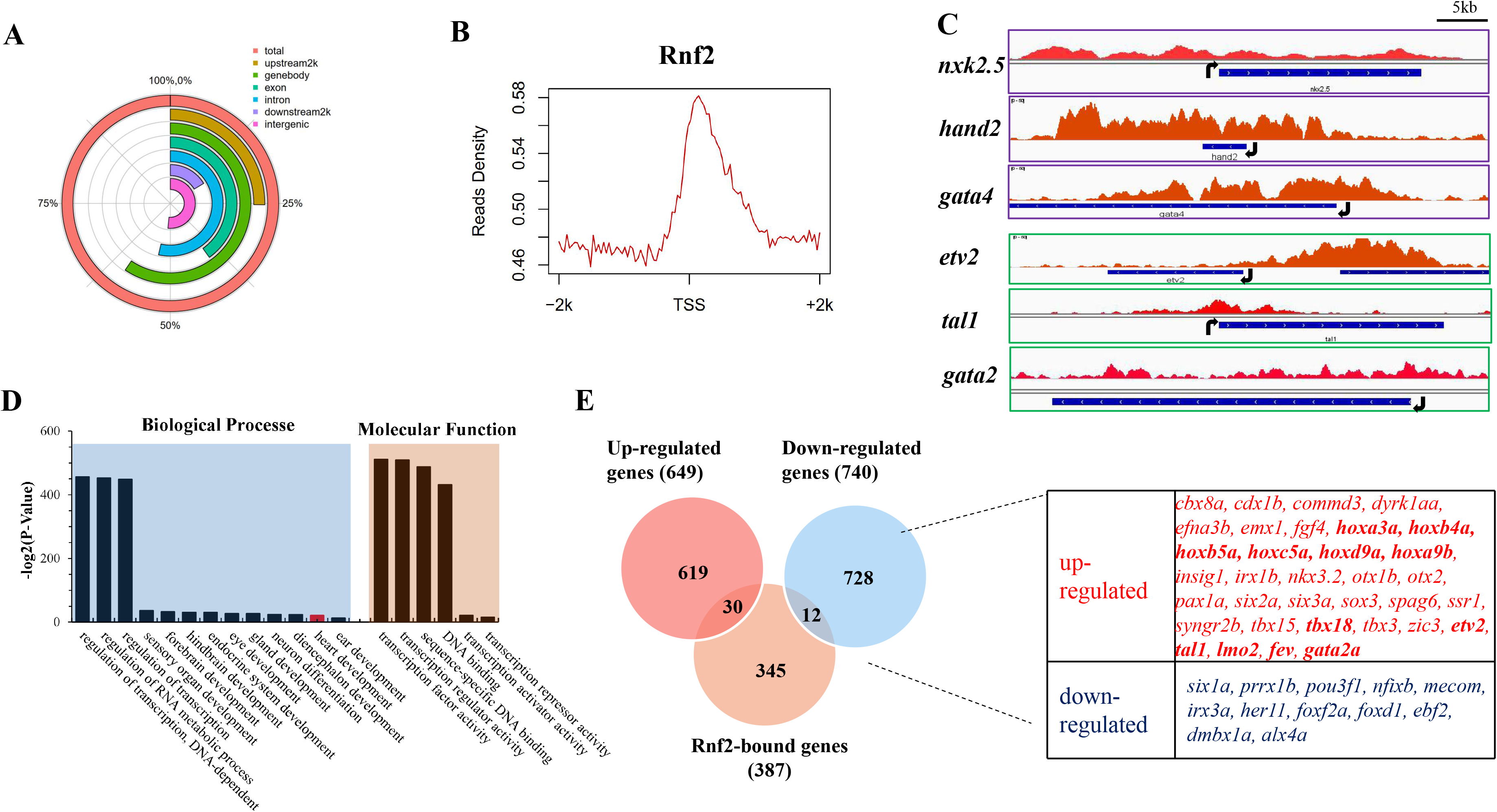
Genome-wide binding profile of Rnf2. **A** Multiple donuts chart showing the binding regions of Rnf2. **B** ChIP-seq reads density of Rnf2, showing Rnf2 peaks at the TSS region. **C** Genome browser snapshots of Rnf2 occupancy at *nkx2.5*, *hand2*, *gata4*, *etv2*, *gata2* and *tal1*. Black arrows showing the transcription direction. **D** GO analysis of Rnf2 target genes. **E** Venn diagram showing the numbers of up- and down-regulated genes that are bound by Rnf2. Dashed line linked boxes showing the up- and down-regulated Rnf2 bound TF genes. The Rnf2 ChIP-seq experiments were repeated two times.

**Table 3.**
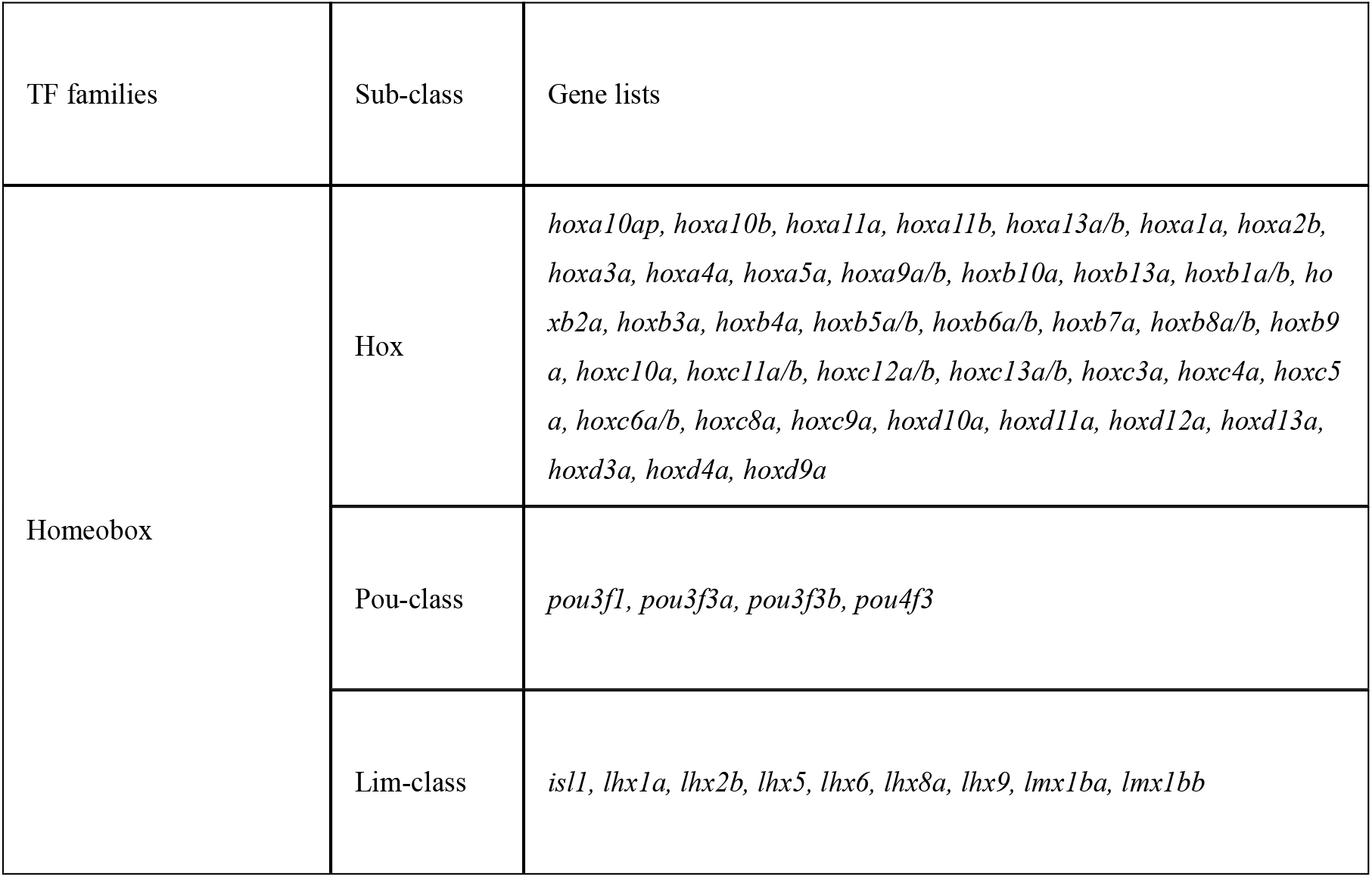
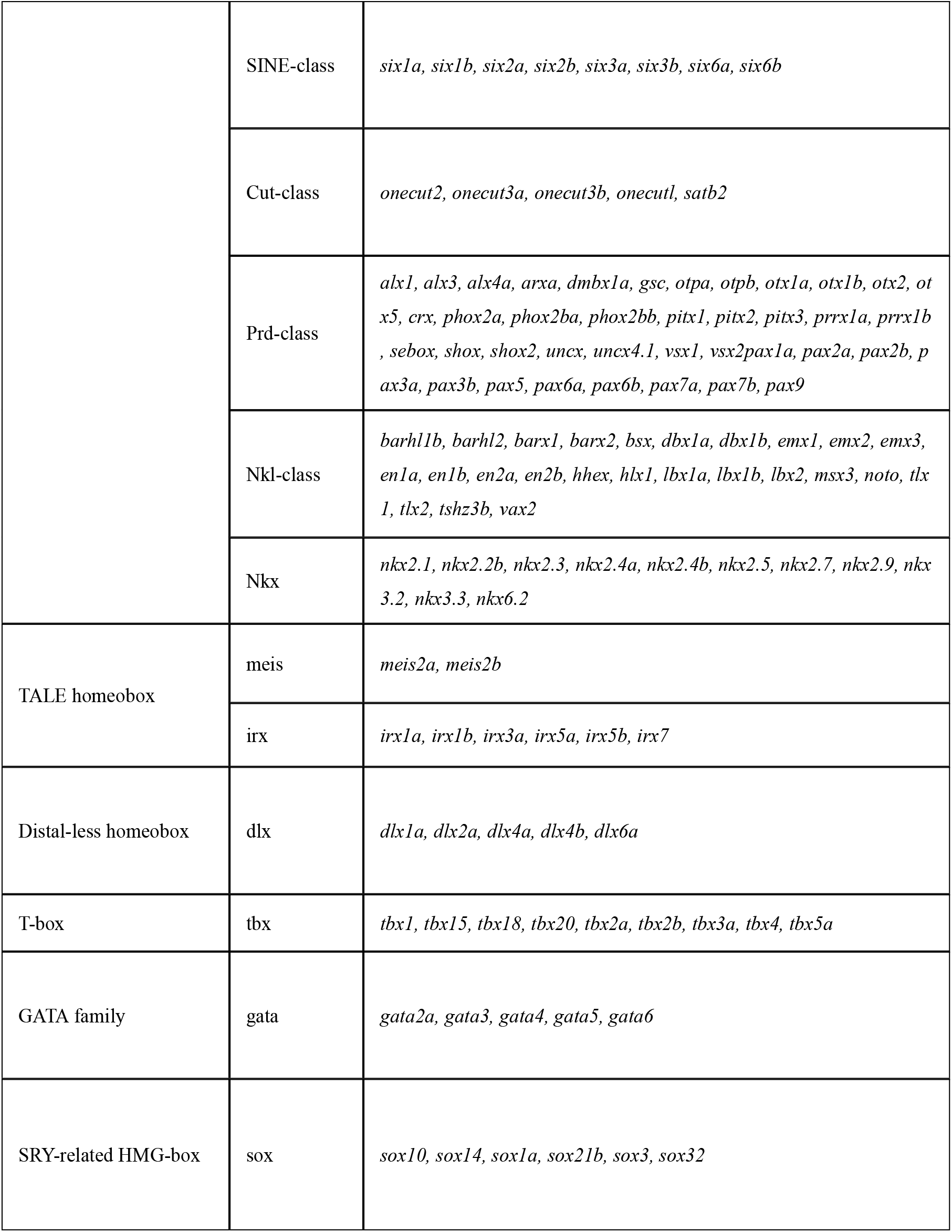
Rnf2-bound family of transcription factors

The combined analysis of ChIP-seq and RNA-seq data allowed us to gain insights into how Rnf2 regulates cardiac and hematoendothelial gene expression. Surprisingly, we found that the majority (approximately 90%) of Rnf2-bound genes remained unchanged by loss of Rnf2. For instance, at 24 hpf, a total of 42 Rnf2-bound genes were deregulated, representing only ∼10% of all Rnf2-bound genes (Fig. 7E). Of the 42 genes, 30 were up-regulated and 12 were down-regulated, which was in line with that Rnf2 functions primarily as a transcriptional repressor.

Next, we grouped Rnf2-bound developmental regulatory genes into two clusters according to their expression state in the hearts of 24 hpf wild-type embryos. Cluster 1 genes are normally expressed in cardiovascular lineages, including early cardiac specification genes *isl1*, *hand2*, *nkx2.5* and *gata4* as well as hematoendothelial progenitor genes such as *nfatc1* and *etv2*; and cluster 2 consisted of genes which are normally not expressed or lowly expressed, including neural genes such as *sox10*, *dlx5* and *pax6* as well as endodermal genes such as *gata6* and *sox17* (Supplementary Fig. 7A-B). We found that cluster 2 genes remained undetectable or lowly expressed in the mutant hearts, suggesting that they remained to be silenced independent of Rnf2. Cluster 1 genes exhibited different scenario in the absence of Rnf2. The hematoendothelial progenitor genes such as *gata2*, *tal1, hox4b, etv2* and *nfatc1* were significantly up-regulated, whereas early cardiac specification genes such as *gata4/isl1/nkx2.5* were barely de-regulated. These observations indicated that the expression of endocardial/hematoendothelial progenitor genes appears more sensitive to loss of Rnf2 function than early cardiac regulatory genes (see below for mechanism).

The chamber myofibril and calcium handling genes, which were significantly down-regulated in the absence of Rnf2, are largely devoid of Rnf2 binding (Supplementary Fig. 7A). This observation further supports that myocardial genes are not directly regulated by Rnf2, and down-regulation of these genes is likely due to the increased expression of hematoendothelial progenitor genes.

### Hematoendothelial progenitor genes are suppressed by Rnf2 via its H2Aub1 catalytic activity

RNF2 is the core enzyme of the PRC1 complex that catalyzes histone H2A lysine 119 ubiquitylation to generate H2Aub1 mark which is associated with gene repression (Blackledge et al., 2014; King et al., 2018; Tamburri et al., 2020). Recently it has been suggested that gene repression activity of Rnf2 is primarily mediated by H2Aub1 which is deposited primarily by non-canonical forms of PRC1 (Boyle et al., 2020; Fursova et al., 2019). We therefore asked whether Rnf2-mediated repression of hematoendothelial genes depends on its H2Aub1 catalytic activity in zebrafish. To this end, we performed rescue experiments by injecting mRNAs encoding a mutated version of Rnf2 (Rnf2 I55A) or encoding wild-type Rnf2 proteins into one-cell stage of *rnf2−/−* embryos. The *rnf2 I55A* mutant (Rnf2m) is analogous to the previously described *Rnf2 I53A* mutant in mice, which preserves PRC1 assembly but results in loss of H2AK119ubi (Illingworth et al., 2015) (Fig. 8A; Supplementary Fig. 8A). We confirmed that the *rnf2 I55A* mutant was indeed expressed similar to wild-type levels but lacked the H2A ubiquitination activity (Fig. 8B). Injection of wild-type *rnf2* mRNAs partially rescued the expression of selected cardiac chamber genes as well as the pleiotropic phenotypes (including edema, cardiac morphogenesis as well as the hematopoietic defects) (Fig. 8C; Supplementary Fig. 8B). However, injection of different dose of *rnf2 I55A* mRNAs had no obvious effects. The rescue results suggested that the catalytic activity of Rnf2 (therefore the H2Aub1 histone modification) is important for repressing key hematoendothelial progenitor genes, which is important for cardiovasculogenesis and hematopoiesis.

**Fig. 8.**
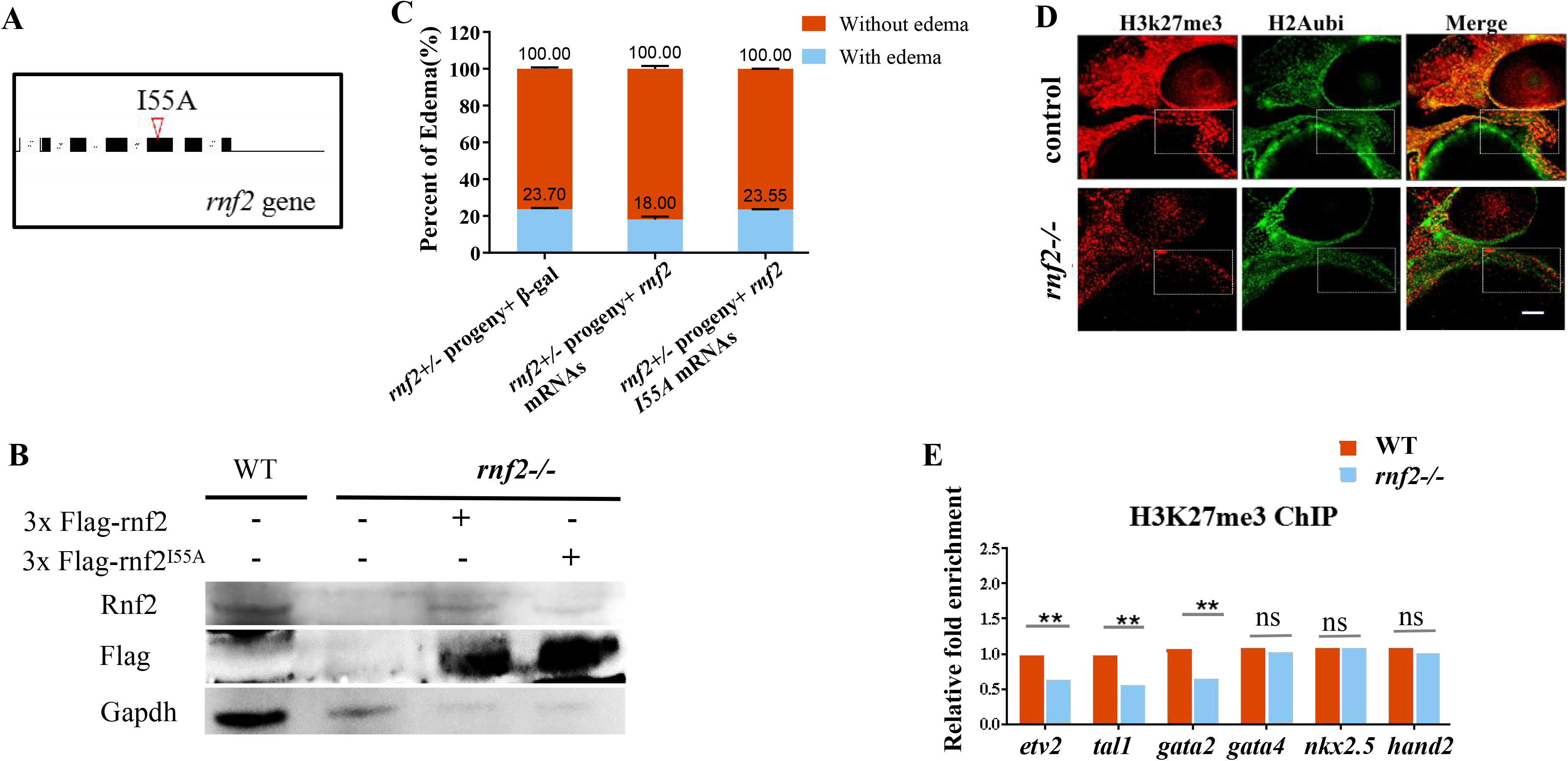
Rnf2 represses hematoendothelial genes via depositing H2Aub1. **A** Cartoon showing the generation of the *rnf2 I55A* mutant. **B** WB showing the expression of Rnf2 I55A protein that is unable to generate H2Aub1. 3x *Flag-rnf2* or *3x Flag-rnf2 I55A* mRNAs were injected into *rnf2* mutant embryos, and lysates were made for detecting the indicated proteins. **C** Percentage of edema of *rnf2+/−* incross progeny that were injected with *β-gal*-, *rnf2*- and *rnf2* I55A- mRNAs. **D** Representative images of cardiac regions by double IF staining using H3K27me3 and H2Aub1 antibodies. White bar: 0.2 mm. **E** ChIP-PCR analysis of hematoendothelial and cardiac specification genes using H3K27me3 antibodies using isolated hearts from *rnf2* mutant and control embryos. **, *P≤* 0.01; ns: not significant.

It has been shown that PRC1-mediated H2Aub1 is sufficient to recruit PRC2/H3K27me3 in at least some contexts, and vice versa (Blackledge et al., 2014; Rougeot et al., 2019). PRC1/RNF2-deposited H2Aub1 and PRC2/EZH2-catalyzed H3K27me3 reinforce each other’s binding to silence the developmental genes (Fursova et al., 2019), and loss of H2Aub1 and/or H3K27me3 leads to de-repression of developmental genes (Sun et al., 2020). We performed immunofluorescence (IF) staining for 24 hpf control and mutant embryos using H2Aub1 and H3K27me3 antibodies. As expected, H2Aub1 levels were greatly reduced in mutant hearts (Fig. 8D). A global reduction of H3K27me3 levels was also observed, which was in line with that H3K27me3 deposition is dependent on H2Aub1 levels (Tamburri et al., 2020).

To investigate whether H3K27me3 mark was affected at individual hematoendothelial progenitor genes or early cardiac specifying genes in the absence of Rnf2, we performed ChIP-PCR experiments for control and mutant hearts using H3K27me3 antibodies. The results showed that H3K27me3 levels were reduced to a greater extent at key hematoendothelial progenitor genes than at early cardiac specifying genes (Fig. 8E; Supplementary Fig. 8C). To understand the possible mechanism behind this, we examined the bivalent domains of these genes. Rnf2/H2Aub1 levels/domains were much lower/narrower at *etv2/tal1/gata2* genes than at *nkx2.5/gata4/hand2* genes (Fig. 7C) (Chrispijn et al., 2019). In mammalian cells, it has been shown that H2Aub1-positive bivalent promoters more efficiently retain the repressive H3K27me3 upon differentiation than PRC1-negative counterparts (Ku et al., 2008), suggesting that H2Aub1 levels negatively affect gene activation of developmental genes during cell lineage restriction. This may interpret why a greater reduction of H3K27me3 levels was observed in hematoendothelial progenitor genes than early cardiac specifying genes in the absence of Rnf2. As H3K27me3 is associated with gene repression, it also may interpret why the expression of hematoendothelial progenitor genes were more sensitive to loss of Rnf2 than early cardiac specifying genes, although this requires further validation.

### Sarcomere assembly and calcium handling were defective in *rnf2* mutants

Since many myofibril- and sarcomere-related genes were de-regulated in the absence of Rnf2, we hypothesized that the cardiac sarcomere formation might be defective in *rnf2−/−* embryos. To this end, we examined the cardiac structure of 80 hpf control and mutant embryos by using the transmission electron microscope (TEM). First, the mutant hearts were obviously hypotrophy compared to the controls, which was consistent with the reduced expression of profibrotic genes. The sarcomere is the basic structural and functional unit of the fibril. It is bordered by a Z band on each end with adjacent I bands, and there is a central M-line with adjacent H-bands and partially overlapping A-bands. The TEM results showed that *rnf2−/−* embryos grossly had a sarcomere structure similar to the control embryos (Fig. 9A). However, there were defects in sarcomere alignment in mutant embryo hearts. First, the myofibril fibers in *rnf2−/−* embryos were arranged more tightly than that in control embryos (Fig 9A). Second, the I-band and Z-disc in *rnf2−/−* embryos were significant wider than that in wild-type embryos (Fig. 9B). Third, the distance between two adjacent Z disks was obviously longer in the hearts of mutants compared to controls, which was in line with that there was a cardiac contraction problem in mutant embryos. Thus, loss of Rnf2 led to a defective cardiac sarcomere alignment, which may contribute to the cardiac contractile defects in the mutant embryos.

**Fig. 9.**
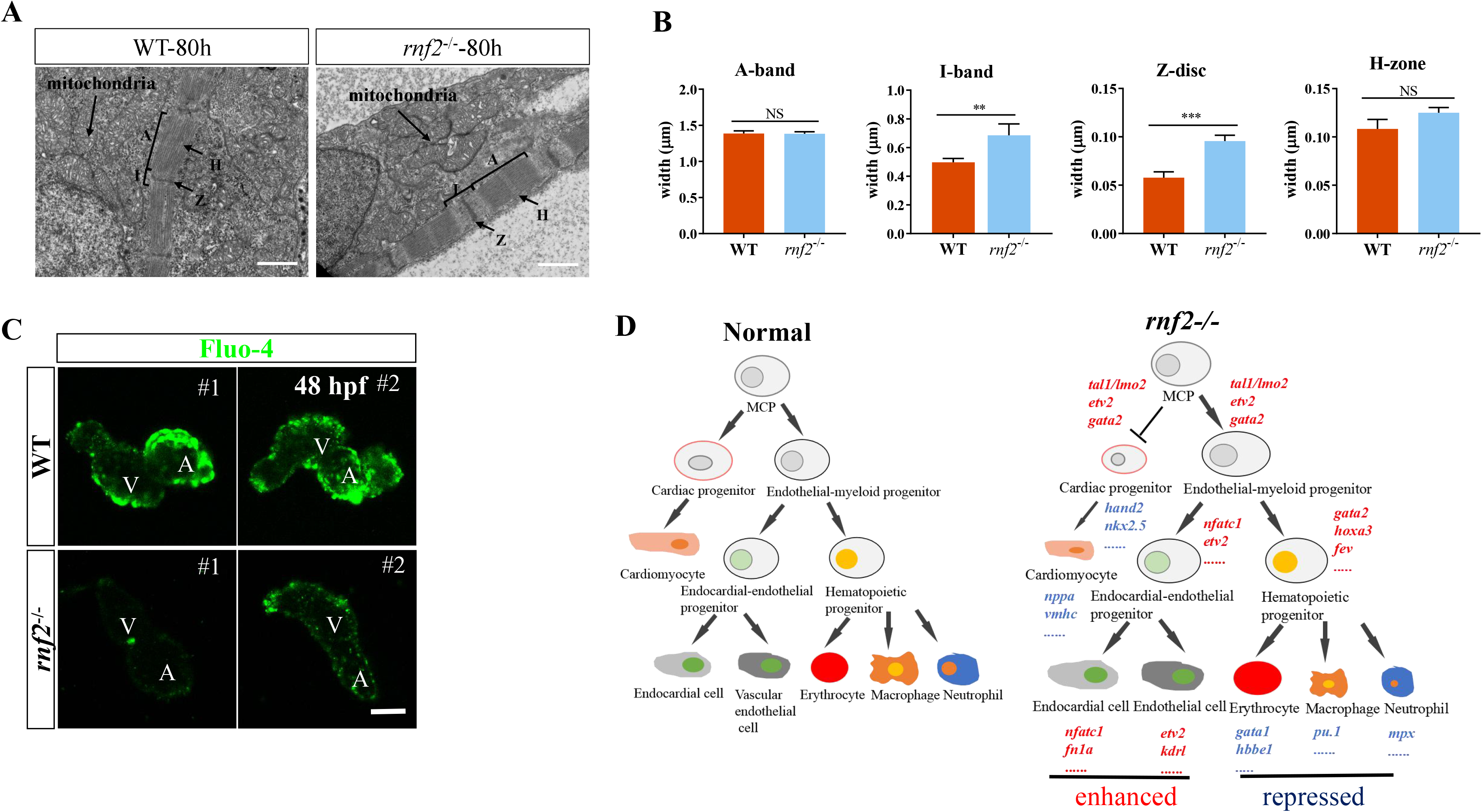
Sarcomere assembly and calcium handling were defective in *rnf2* mutants. **A** Cardiac TEM assay revealed that the sarcomere of cardiac muscle was abnormal in *rnf2−/−* hearts. A, A-band; I, I-band; H, H-zone; Z, Z-disc. Scale bar: 1.0 μm. **B** Bar graph showing the width of A-band, I-band, Z-disc and H-zone in cardiac sarcomere of wild type and *rnf2−/−* embryos in A. **C** Fluo-4 staining showing the Ca2+ concentration in *rnf2−/−* and wild-type hearts at 48 hpf. A, atrium; V, ventricle. **D** Schematic diagram illustrating the mechanism by which *rnf2* regulates the coordinated cardiovascular and hematopoietic lineage commitment and development. *, *p <* 0.05; **, *p <* 0.01; ***, *p <* 0.001; NS, no significant.

As many calcium handling genes were deregulated in hearts of mutant embryos, we investigated whether calcium signaling was defective in mutant embryos. To this end, we detected the calcium signal intensity of single hearts for control and mutant embryos using calcium-sensitive dye, fluo 4-AM. Calcium indicator Fluo 4-AM was widely used to detect the fluorescence-marked calcium flux in heart by confocal imaging (Puceat, 2013). The result showed that the calcium signal intensity was dramatically reduced in *rnf2*^−/−^ embryos compared to the controls (Fig. 9C).

## Discussion

PcG proteins have been shown to stabilize transcriptional programs in embryonic progenitors and their differentiated descendants (Rougeot et al. 2019; Delgado-Olguin et al. 2012). However, the exact mechanisms by which PcGs control embryonic cardiovascular and hematopoietic development remain unclear, likely due to embryonic arrest or lethal of the mice. Zebrafish embryos can survive a week also without an intact circulating system and hence provide an excellent system to study the roles of PRC1/Rnf2 in cardiovascular and hematopoietic development during embryogenesis.

To our knowledge, this is the first report showing that Rnf2, the core enzymatic component of Polycomb repressive complex 1 (PRC1), plays an important role in the control of cardiovascular and hematopoietic development and differentiation via suppressing the master hematoendothelial progenitor genes in zebrafish, and its loss leads to up-regulation of these genes, resulting in severe defects in cardiogenesis and hematopoiesis. The role of PRC1/Rnf2 in suppressing hematoendothelial progenitor genes during vertebrate embryogenesis has not been described before. The repression activity of Rnf2 is persistently required, ranging from early somitogenesis in the ALPM, cell lineage commitment, to organ homeostasis and functional maturation. A remarkable example is that hematoendothelial progenitor genes such as *nfatc1* and *gata2* and hematopoietic-lineage gene such as *rag1* were ectopicaly expressed in 48 hpf mutant hearts, which will likely contribute to the abnormal cardiac gene expression program, consequently, the severe cardiac phenotypes.

By combining both ChIP-seq and RNA-seq data, we show that Rnf2 directly binds to and represses the key hematoendothelial progenitor genes such as *tal1* and *gata2*. We show that Rnf2-mediated repressive mechanism is dependent on its H2Aub1 catalytic activity and may be also involved in recruiting PRC2/H3K27me3. Importantly, injection of *rnf2* mRNAs can simultaneously rescue both the cardiac and hematopoietic phenotypes. Together, our results revealed that PRC1/Rnf2-mediated epigenetic mechanism is important for the precise control of gene expression that is critical for both cell lineage separation and the coordinated cardiovascular and hematopoietic development in zebrafish (Fig. 9D).

The endocardial/hematoendothelial progenitor genes such as Tal1/Lmo2 and Gata2 function at the top of the regulatory cascade that is essential for both cardiovasculogenesis and hematopoiesis (Chen et al., 2012; Palencia-Desai et al., 2011; Schumacher et al., 2013; Schupp et al., 2014). Lmo2, Tal1 and Gata2 form a gene-regulatory circuit during the specification of MCPs (Pimanda et al., 2007; Shi et al., 2014). Gain of function of Tal1/Lmo2/Gata2 leads to an expansion of endothelial and hematopoietic cells at the expense of myocardial precursors, and loss of function studies shows the opposite effect (Castano et al., 2019; Elcheva et al., 2014; Patterson et al., 2007). Etv2 and Nfatc1 are the earliest transcriptional regulators that are expressed in precursors of cardiovascular endothelial cells in both zebrafish and mouse embryos (Puceat, 2013; Saint-Jean et al., 2019; Wong et al., 2012). Transcriptional inhibition of *etv2* is essential for embryonic cardiac development (Schupp et al., 2014), and overexpression of *etv2* results in the reduced expression of endogenous myocardial markers and the expansion of both endocardial and myocardial markers (Palencia-Desai et al., 2011; Sumanas and Lin, 2006). Thus, the proper expression of hematoendothelial progenitor genes is critical for both cardiovascular and hematopoietic development, and therefore need to be tightly regulated by PRC1/Rnf2.

We found that loss of Rnf2 leads to severe defects in both the primitive and definitive wave of hematopoiesis during zebrafish embryogenesis. For the primitive hematopoiesis, the expression of early erythroid markers was increased, while the expression of myeloid- and granulocyte-markers was decreased. During the definitive hematopoiesis, although HSC numbers are increased in the absence of Rnf2, its differentiation into more mature blood cells are blocked, as *rnf2* mutant embryos lack of all three differentiated cell types examined: granulocytes, macrophage and erythrocytes, suggesting that Rnf2 is required for differentiation of blood progenitor cells. Thus, loss of Rnf2 results in increased HSC numbers and arrested differentiation, hallmarks of leukemia. Whether Rnf2 in adult fish has a similar function remains unclear due to the embryonic lethal of *rnf2* mutants. Nevertheless, we anticipate that this is the case (Di Carlo et al., 2018).

In pluripotent cells, developmental genes, including the hematoendothelial regulatory genes and the cardiac specifying genes, are often decorated with both active H3K4me3 mark and repressive H3K27me3/H2Aub1 marks at gene promoters (Ku et al., 2008). The bivalent domains are thought to silence the developmental genes while keeping them poised for later activation. These developmental genes become activated when the repressive histone marks are removed during cell differentiation. Surprisingly, the cardiac specification genes such as *isl1, hand2* and *nkx2.5*, were barely deregulated in the absence of Rnf2, in contrast to key hematoendothelial progenitor genes. It has been shown that H2Aub1-positive bivalent promoters more efficiently retain H3K27me3 upon differentiation than PRC1-negative counterparts, suggesting an important role of H2Aub1 or H2Aub1 levels in control gene expression. We found that Rnf2/H2Aub1 levels/domains are much lower/narrower at *etv2/tal1/gata2* genes than at *nkx2.5/gata4/hand2* genes, and that loss of Rnf2 has a greater effect on H3K27me3 levels at hematoendothelial progenitor genes than at cardiac specification genes. This may partially explain why the hematoendothelial progenitor genes are more sensitive to loss of Rnf2. In addition, recent studies have suggested that RNF2-regulated nucleosome occupancy and spacing as well as high-order genome structure are also important for gene expression state (Fursova et al., 2019; King et al., 2018). Whether these mechanisms contribute to gene expression regulation awaits future investigation.

Chrispijn et al. (2019) has reported that loss of Rnf2 leads to up-regulation of a few direct target genes such as *tbx2* and *tbx3* (bound by Rnf2) which are negative regulators of cardiac chamber and structural genes, resulting in defects in cardiac looping and function. Unfortunately, the role of hematoendothelial progenitor genes has not been discussed in detail although the up-regulation of *tal1* was shown by a RNA-seq snapshot in that work (Chrispijn et al., 2019). Compared to Chrispijn, in our hand, a significant larger number of DEGs (a total of more than 1,000 genes) were observed in single hearts of 24 hpf mutant embryos. The discrepancy might be due to the different RNA-sequencing depth between us and Chrispijn et al. In our hand, an average of 40-50 M clean reads were obtained per heart, and approximately 18,000 genes were detected. The high RNA-sequencing depth may allow us to detect more DEGs by loss of Rnf2 function. Because the majority of the down-regulated DEGs, particular the chamber genes, are devoid of Rnf2 binding, we propose that a direct role of Rnf2 (Rnf2-bound genes such as *tbx2/3* genes) and an indirect role of Rnf2 (secondary to the up-regulation of hematoendothelial TF genes) are present to control cardiac gene expression.

## Acknowledgement

We thank Prof. Yiyue Zhang from South China University of Technology for the critical comments of the manuscript, and Prof. Donghui Zhang from Hubei University for the help with calcium signaling analysis. This work was supported by the Strategic Priority Research Program of Chinese Academy of Sciences (Grant No. XDB31000000); National Science Foundation of China (NSFC #31671526), and National Key Research and Development Program of China (2016YFA0101100).

## Legend for supplementary figures

**Fig. S1.**
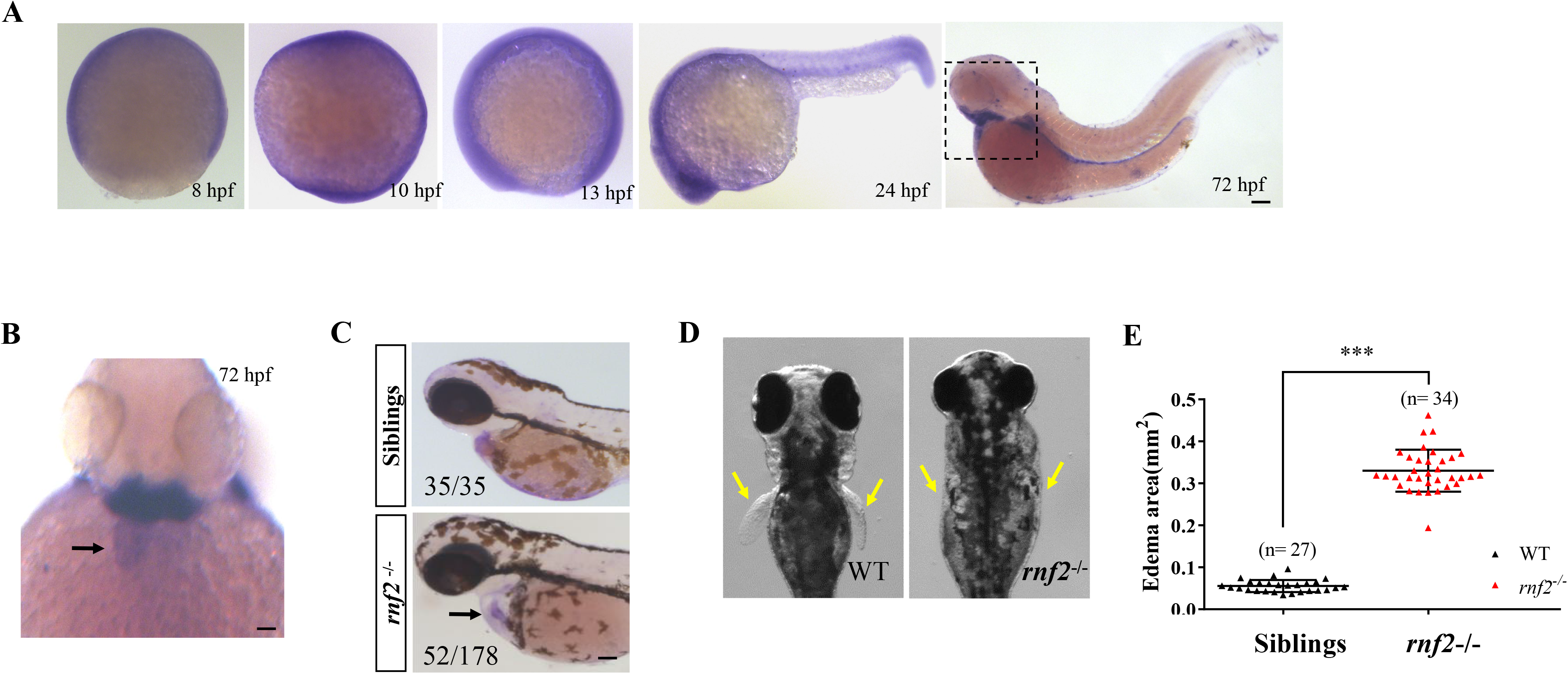
**A** Whole mount in situ hybridization (WISH) of *rnf2* at the indicated time points. **B** Close-up of the dashed box in A. Arrow showing that *rnf2* was expressed in the heart. Ventral view. **C** In situ hybridization of *cmlc2.* Black arrow showing the *stringy heart* phenotype in the mutant embryo. **D** Yellow arrows showing the absence of pectoral fins in mutant embryos. **E** Edema area measurement for control and mutant embryos at 3 dpf. *** indicates *p<* 0.001.

**Fig. S2.**
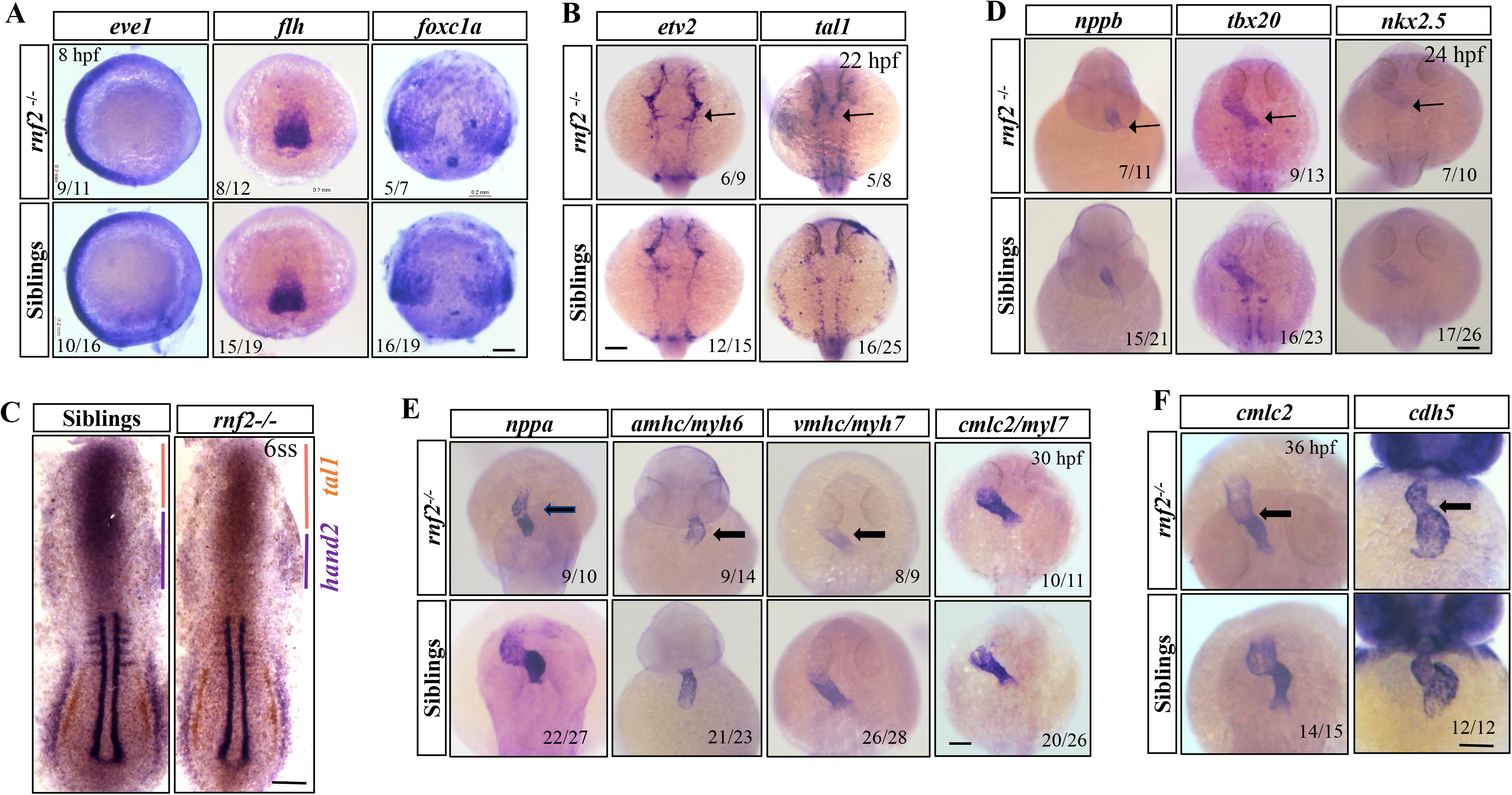
**A** WISH results of control and mutant embryos at 7 hpf, showing the expression of early mesodermal markers *eve1*, *flh* and *foxc1a*. Scale bar: 0.2 mm. **B** WISH results of control and mutant embryos at 22 hpf, showing the expression of *ev2* and *tal1*. Scale bar: 0.2 mm. **C** WISH results for *tal1/hand2/myoD*. In the ALPM, *tal1* and *hand2* are expressed in proximity. Note that there was a slight expansion of *tal1* (orange) and a slight reduction of *hand2* (purple) in the mutant embryo compared to controls. *MyoD* staining was used for determination of the precise developmental stage of embryos. Anterior is upwards. **D** WISH results showing the reduced expression of cardiac genes *nppb*, *nkx2.5* and *tbx20* in mutant embryos compared to the controls at 24 hpf. Scale bar: 0.2 mm. **E** WISH results showing the reduced expression of cardiac genes *nppa*, *amhc*, *vmhc* and *cmlc2* in mutant embryos compared to the controls at 30 hpf. Scale bar: 0.2 mm. **F** WISH results of control and mutant embryos at 36 hpf, showing the expression pattern of *cdh5* and *cmlc2*. Note the cardiac looping (black arrow) and morphogenesis defects in mutant embryos. Scale bar: 0.2 mm.

**Fig. S3.**
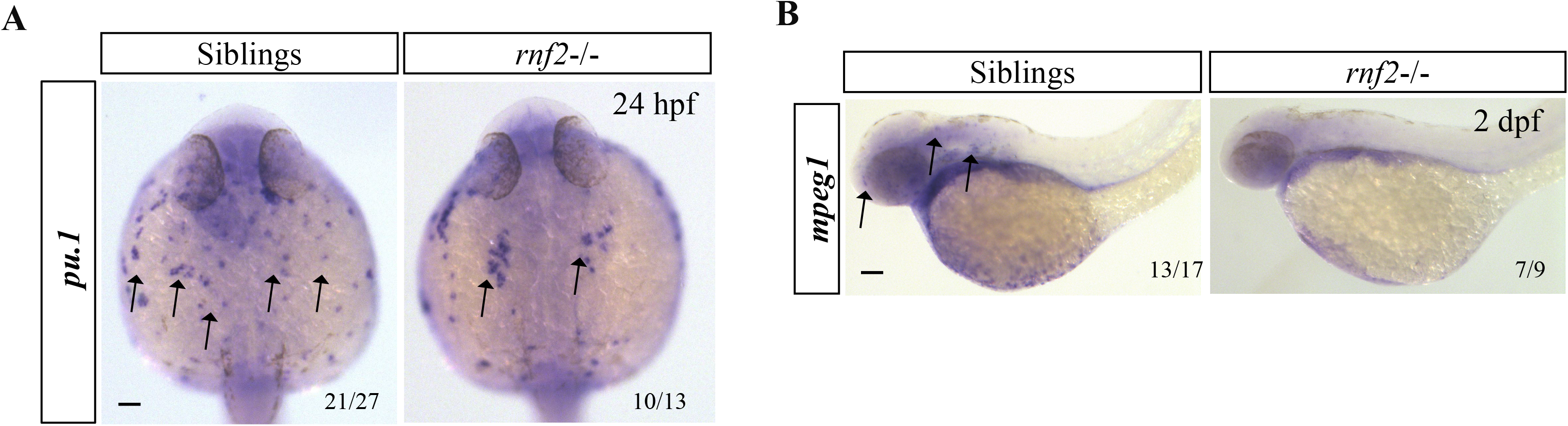
**A** WISH results for *pu.1*, showing the reduced signals in yolk of mutant embryos compared to the controls. Ventral view. **B** WISH results for the macrophage marker *mpeg1*, showing the reduced *mpeg1* positive signals in head regions of mutant embryos compared to the controls. All experiments were repeated three times and images are representatives.

**Fig. S4.**
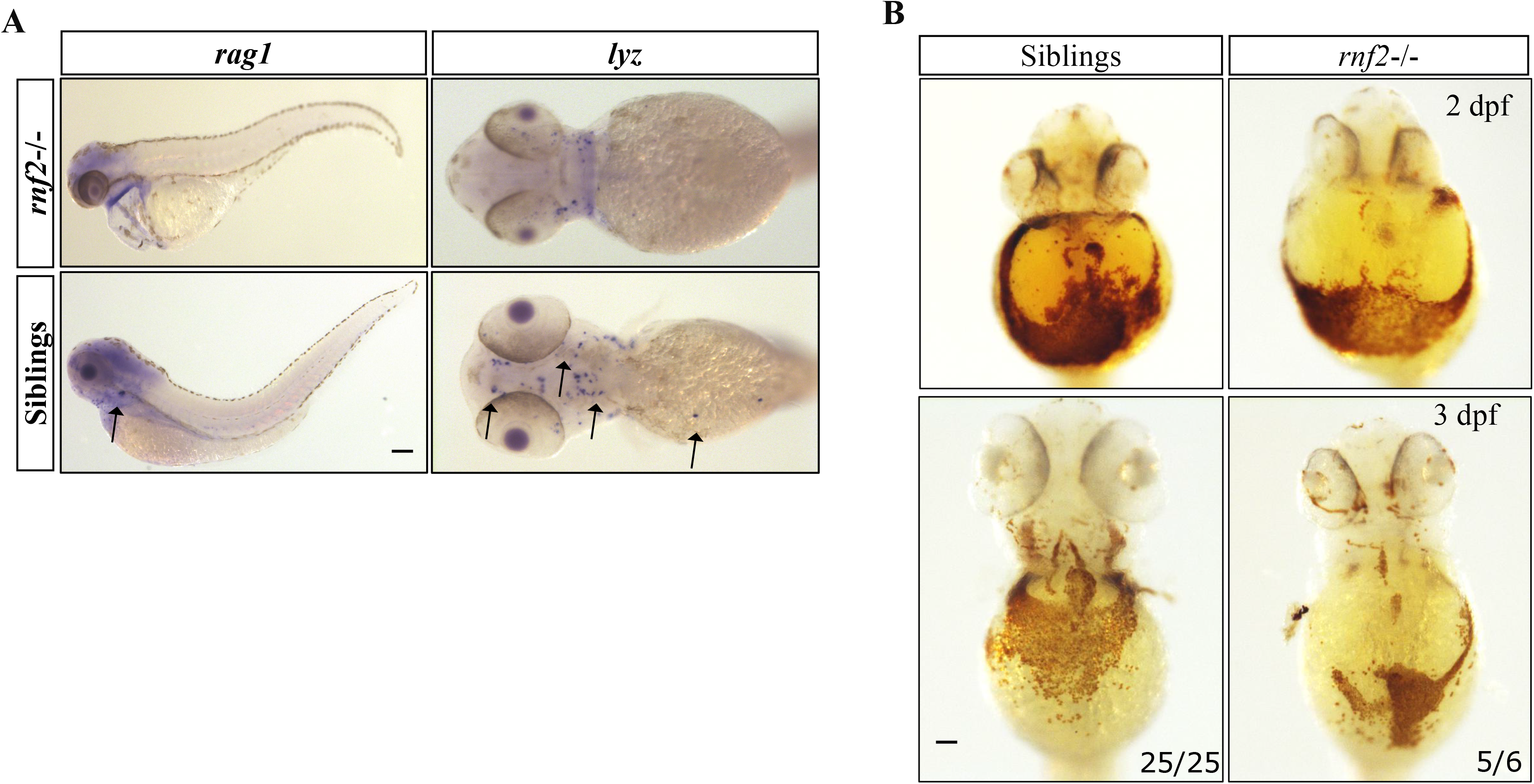
**A** WISH results for *rag1* and *lyz* in control and *rnf2* mutant embryos. Left: Black arrow pointing to the *rag1*-expressing thymus in control embryos, while the signal was not visible in mutant embryos. Lateral view. Right: ventral view showing the reduced *lyz* positive signals in mutant embryos in head and yolk sac regions. **B** Ventral view of O-Dianisidine staining of 2 or 3 dpf control and mutant embryos. Ventral view for head and yolk sac regions.

**Fig. S5.**
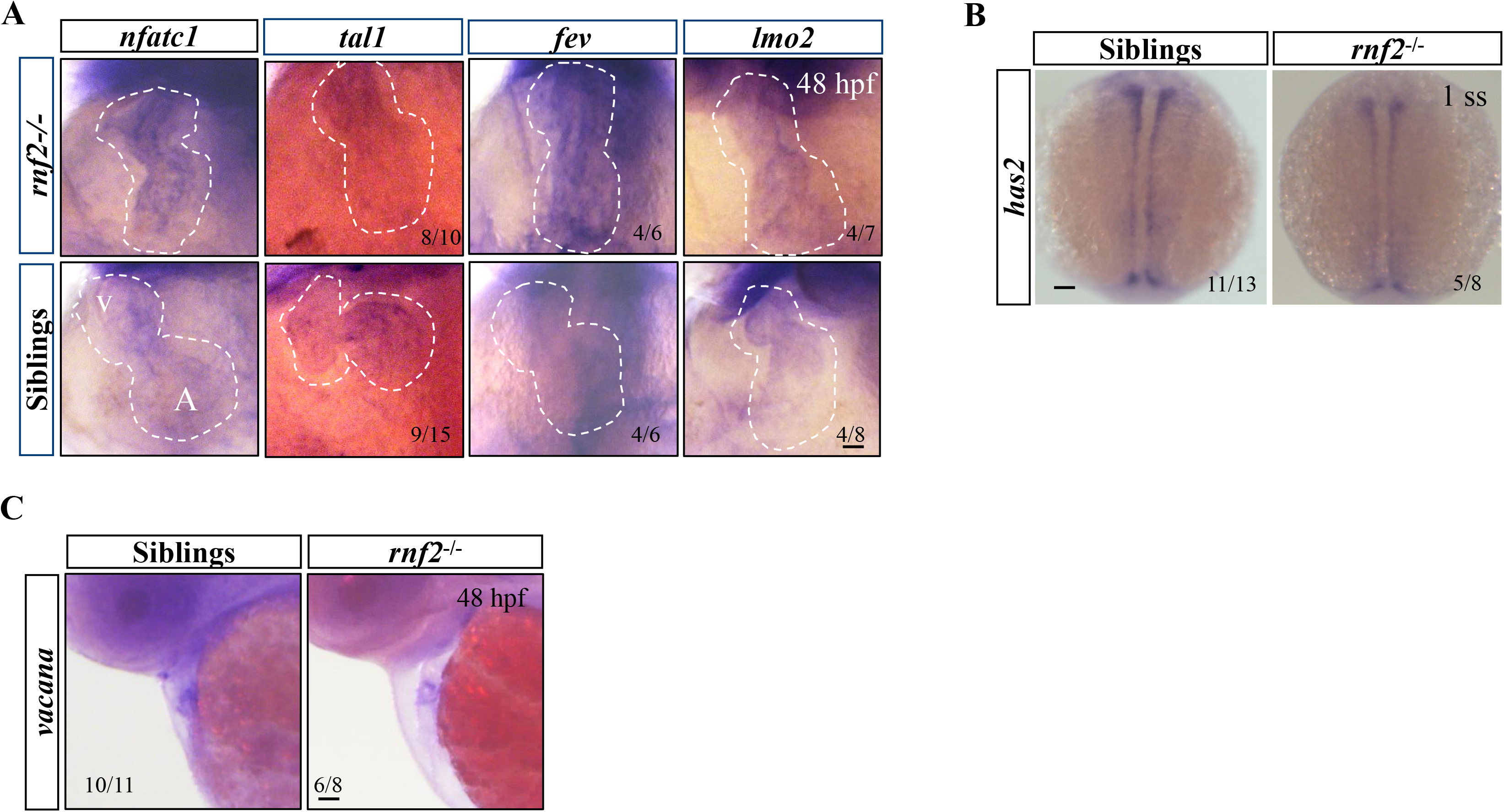
**A** WISH results for control and mutant embryos, showing the expression of the indicated markers. V: ventricular; A: atrial. **B** WISH results showing the expression of *has2* in 1-somite stage control and mutant embryos. **C** WISH results showing the reduced expression of the AVC marker *vacana* in mutant hearts compared to controls. Lateral view. All experiments were repeated three times and images are representatives.

**Fig. S6.**
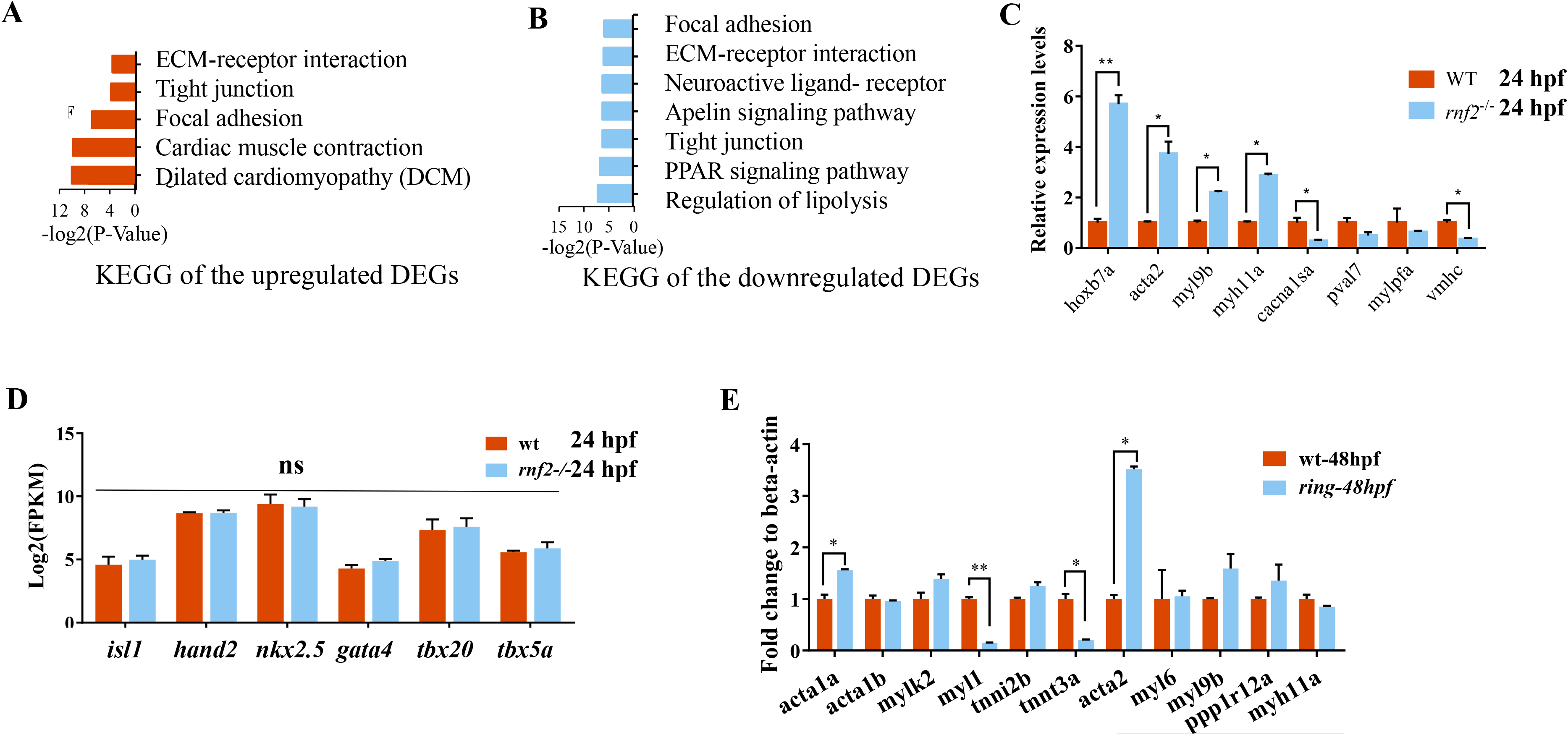
RNA-seq analysis of 24 hpf embryos. **A** KEGG analysis of the up-regulated DEGs from whole control and mutant embryos at 24 hpf. **B** KEGG analysis of the down-regulated DEGs from whole control and mutant embryos at 24 hpf. **C** qRT-PCR analysis of selected cardiac chamber- and muscle-related genes in panel 6C. **D** qRT-PCR analysis of the selected cardiac specification genes, showing no significant change in gene expression in 24 hpf *rnf2* mutant embryos compared to the controls. **E** qRT-PCR analysis of the selected cardiac chamber genes in panel 6G using 48 hpf control and mutant hearts. ns: not significant. All qRT-PCR experiments were repeated three times.

**Fig. S7.**
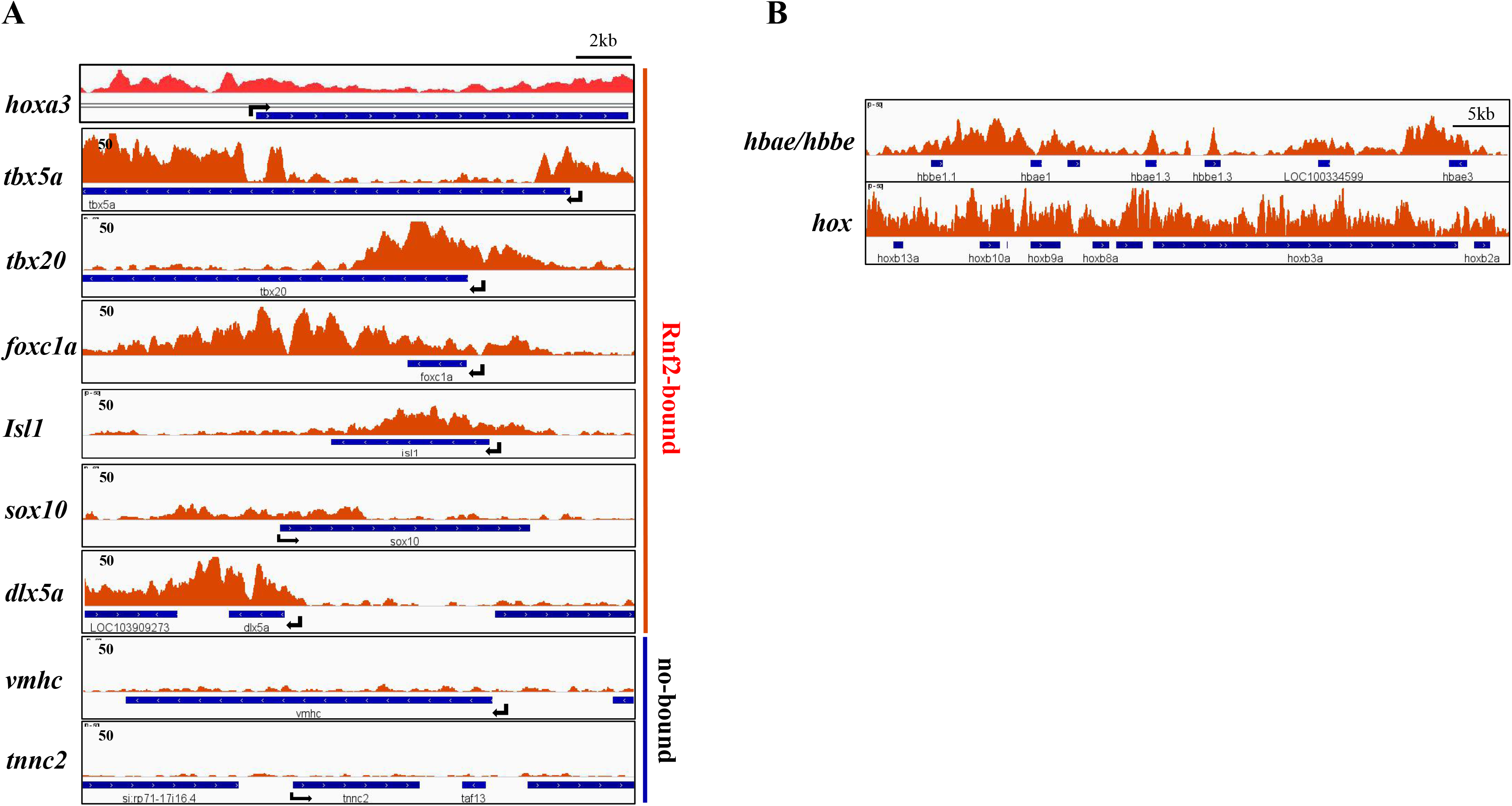
**A** Genome browser snapshots of Rnf2 occupancy at the indicated genes. **B** Genome browser snapshots of Rnf2 occupancy at *hbae/hbbe* and *hox* loci. Black arrows showing the transcription direction.

**Fig. S8.**
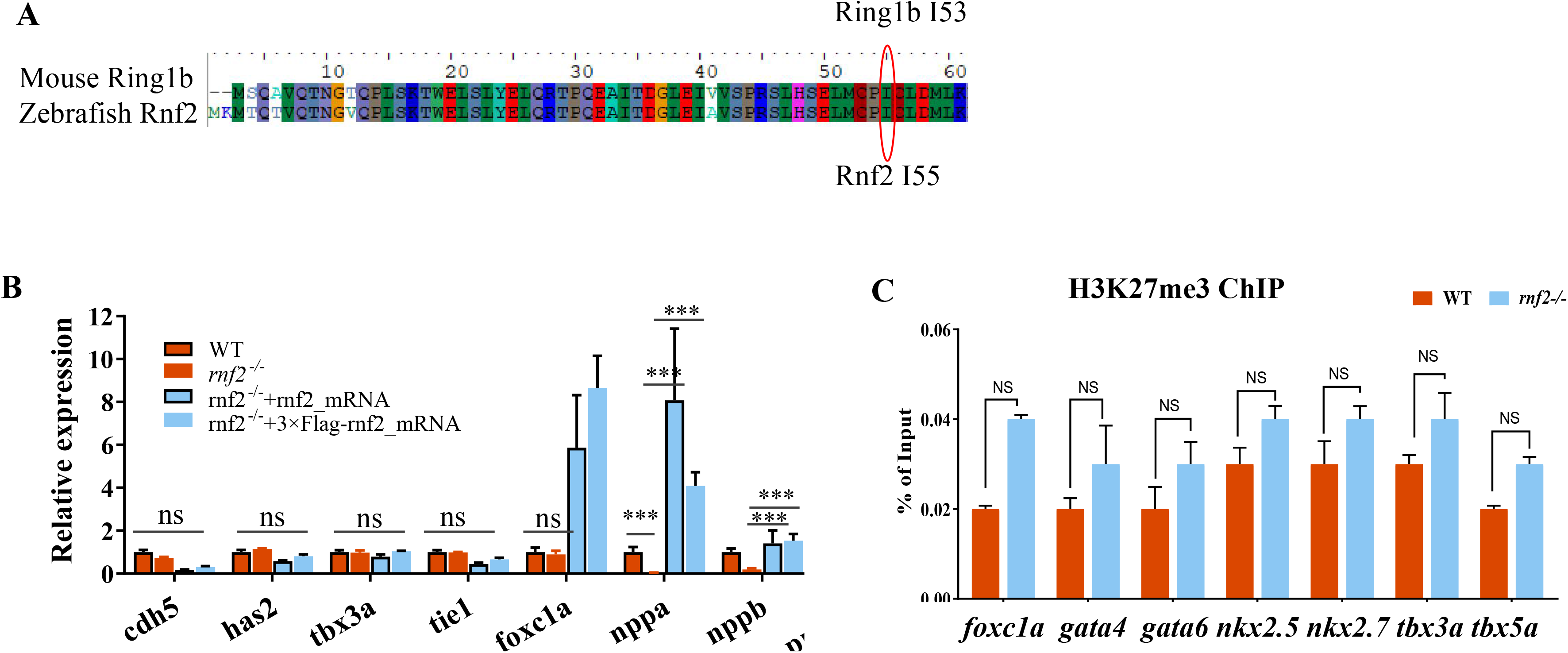
**A** Alignment of the first 60 amino acid residues of mouse Rin1b and zebrafish Rnf2. Note that zebrafish I55 is homologous to mouse I53. **B** H3k27me3 ChIP-PCR analysis for the indicated cardiac genes, showing no change of H3k27me3 levels in the absence of Rnf2. **C** Rescue of expression levels of cardiac chamber genes *nppa* and *nppb* which were down-regulated in *rnf2* mutant hearts by *rnf2* mRNAs. *** indicates *p<* 0.001; ns: not significant. Experiments were repeated two times.

